# Calretinin-expressing islet cells: a source of pre- and post-synaptic inhibition of non-peptidergic nociceptor input to the mouse spinal cord

**DOI:** 10.1101/2023.06.01.543241

**Authors:** Olivia C. Davis, Allen C. Dickie, Marami B. Mustapa, Kieran A. Boyle, Tyler J. Browne, Mark A. Gradwell, Kelly M. Smith, Erika Polgár, Andrew M. Bell, Éva Kókai, Masahiko Watanabe, Hendrik Wildner, Hanns Ulrich Zeilhofer, David D. Ginty, Robert J. Callister, Brett A. Graham, Andrew J. Todd, David I. Hughes

## Abstract

Unmyelinated non-peptidergic nociceptors (NP afferents) arborise in lamina II of the spinal cord and receive GABAergic axoaxonic synapses, which mediate presynaptic inhibition. However, until now the source of this axoaxonic synaptic input was not known. Here we provide evidence that it originates from a population of inhibitory calretinin-expressing interneurons (iCRs), which correspond to lamina II islet cells. The NP afferents can be assigned to 3 functionally distinct classes (NP1-3). NP1 afferents have been implicated in pathological pain states, while NP2 and NP3 afferents also function as pruritoceptors. Our findings suggest that all 3 of these afferent types innervate iCRs and receive axoaxonic synapses from them, providing feedback inhibition of NP input. The iCRs also form axodendritic synapses, and their targets include cells that are themselves innervated by the NP afferents, thus allowing for feedforward inhibition. The iCRs are therefore ideally placed to control the input from non-peptidergic nociceptors and pruritoceptors to other dorsal horn neurons, and thus represent a potential therapeutic target for the treatment of chronic pain and itch.

## INTRODUCTION

The spinal dorsal horn is innervated by primary afferents, with different populations terminating in a lamina-specific pattern^1^. Unmyelinated (C) afferents, most of which function as nociceptors, arborise in the superficial dorsal horn (SDH, laminae I-II). Early studies identified two main classes of nociceptive C fibres, which are commonly referred to as peptidergic and non-peptidergic. These differed in their dependence on growth factors, their termination zone within the dorsal horn, and their ultrastructural appearance^2–4^. The non-peptidergic nociceptors arborise mainly in lamina II, and are believed to form the central axons in what Ribeiro-da-Silva and Coimbra defined as type I synaptic glomeruli^4, 5^. Glomerular central axons receive axoaxonic and dendroaxonic synapses from GABAergic interneurons^4, 6^, and these are thought to underlie presynaptic inhibition of the afferents^7^. In contrast, peptidergic nociceptors, which can be identified by the expression of calcitonin gene-related peptide (CGRP), terminate mainly in laminae I and IIo and typically form simple synaptic arrangements that lack axoaxonic or dendroaxonic synapses^3, 8^. Recent transcriptomic studies^9–12^ have further divided the non-peptidergic (NP) nociceptors into 3 major classes (NP1-3)^9^, defined by expression of the mas-related G protein-coupled receptors MrgD (NP1) or MrgA3/MrgB4 (NP2), or of somatostatin (SST; NP3). Although it is now clear that some of these afferents do express neuropeptides^9, 10^, we use the NP1-3 nomenclature for convenience.

The SDH contains a large number of densely packed interneurons^1, 13^. The majority (∼75%) of these are excitatory glutamatergic cells, while the remaining 25% are inhibitory, and use GABA and/or glycine as their principal fast transmitter^14, 15^. Each of these major types of interneuron can be subdivided into distinct populations, based on morphological, electrophysiological and neurochemical/transcriptomic criteria^16–20^. We have reported that the inhibitory interneurons in laminae I-II can be assigned to 5 largely non-overlapping neurochemical classes, based on expression of parvalbumin (PV), dynorphin and galanin, neuronal nitric oxide synthase (nNOS), neuropeptide Y (NPY) or calretinin (CR)^21^. This finding is consistent with results from studies that have used single-cell/nucleus RNA sequencing to define neuronal populations^18, 19^.

We have shown that PV-expressing cells give rise to axoaxonic synapses on myelinated low-threshold mechanoreceptors (A-LTMRs)^22^, and these are thought to gate tactile input to nociceptive circuits through both presynaptic and postsynaptic inhibitory mechanisms, thus preventing mechanical allodynia^23–25^. The dynorphin/galanin population has been implicated in the suppression of mechanical pain and pruritogen-evoked itch^26–29^, while activating the nNOS cells leads to a reduction in nocifensive reflexes^26^. There is controversy concerning the roles of NPY-expressing inhibitory interneurons. Although initial studies reported that these cells were responsible for gating mechanical itch^30–32^, we have found that they have a much broader role, including the suppression of pruritogen-evoked itch, acute nocifensive reflexes and hypersensitivity in both neuropathic and inflammatory pain models^33^. Relatively little is known about the inhibitory calretinin cells (iCRs), partly because targeting them has proved difficult as they are greatly outnumbered by calretinin-expressing excitatory interneurons^18, 34, 35^. Two transcriptomic studies have identified populations of iCRs in the dorsal horn, with Häring et al^18^ and Sathyamurthy et al^19^ assigning these cells to their Gaba8 and Gaba9, and DI-1 and DI-5 classes, respectively. We have shown that the iCRs in lamina II have dendrites that are rostrocaudally elongated with a restricted dorsoventral extent^34, 36^, corresponding to a morphological class known as islet cells^17^. Lamina II islet cells have axons that arborise extensively within this lamina^16, 17, 37–39^ raising the possibility that they give rise to the axoaxonic synapses on non-peptidergic nociceptors. Here we have used a range of mouse genetic lines to further characterise the iCRs and to test the hypothesis that they are the source of axoaxonic synaptic input to these nociceptors.

## RESULTS

### Anatomical analysis of patched calretinin cells

In order to investigate the axonal arbors of iCRs, we carried out patch-clamp recordings in sagittal slices prepared from BAC transgenic (CR::GFP) mice in which GFP is expressed under control of the calretinin promoter^40^. Recordings were performed as described previously, using Neurobiotin-filled pipettes^34, 36^. We identified 5 cells that showed tonic firing during injection of depolarising current, a characteristic feature of inhibitory calretinin cells^34, 36^. Following completion of recording, the slices containing these cells were processed to reveal the Neurobiotin label with a peroxidase/diaminobenzidine method, which permits combined light and electron microscopic examination. All 5 cells had typical islet morphology, with rostrocaudally-elongated dendrites that gave rise to dendritic spines, and axons that had numerous varicosities and remained in lamina II^17^ (Fig 1a-d). Although preservation of the tissue structure was compromised at the electron microscope (EM) level, we were able to recognise several cases in which axonal boutons belonging to the labelled cells formed axoaxonic synapses on the central terminals of type I synaptic glomeruli^4^. These could be identified by their indented contour and the presence of synaptic vesicles of varying sizes (Fig 1e,f). We also observed synapses at which dendrites of the recorded cells were postsynaptic to the central axons of type I glomeruli (Fig 1g).

**Fig 1.**
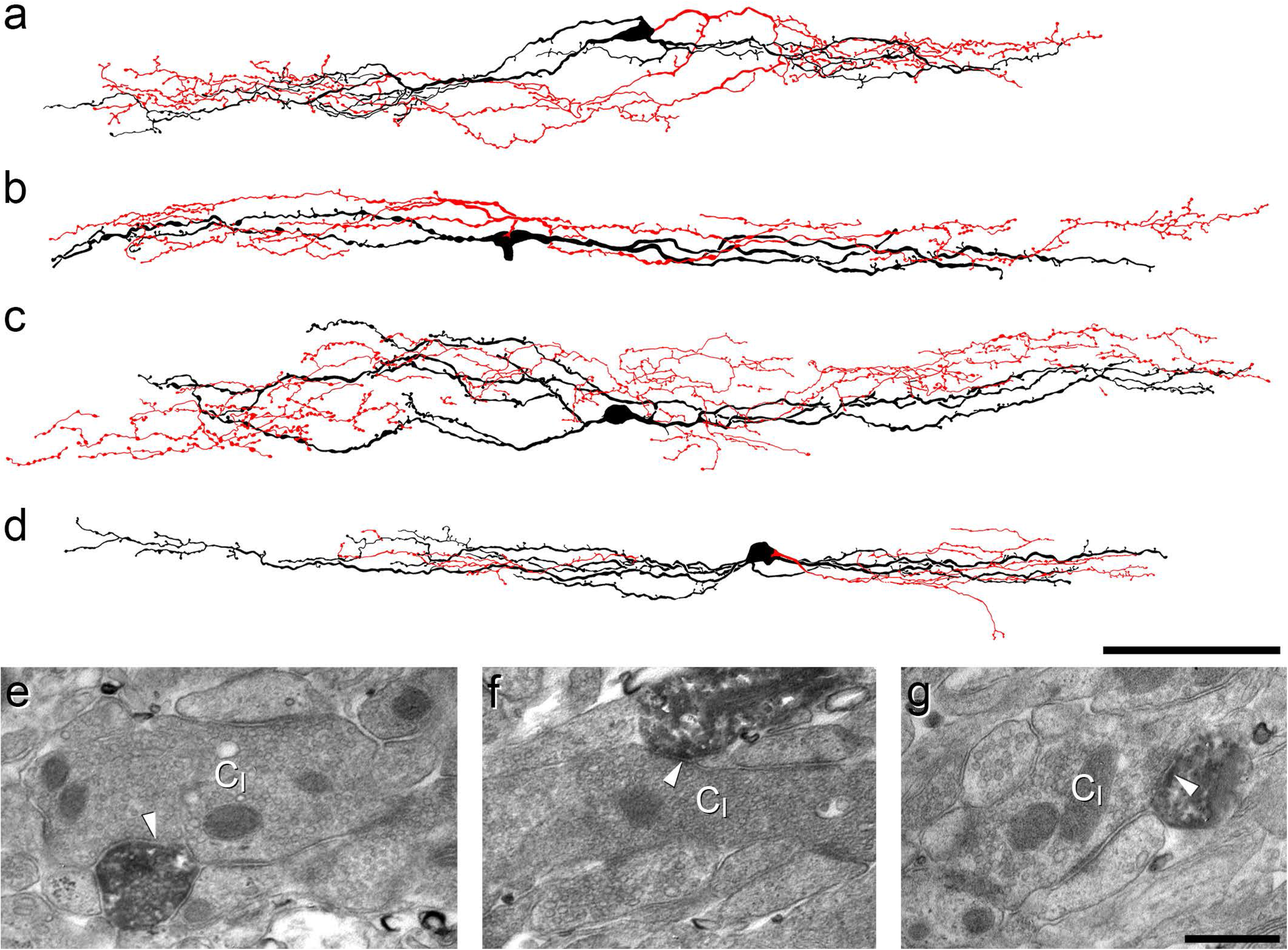
Reconstruction of iCRs and examples of their ultrastructure. **a**-**d** show reconstructions of the soma and dendritic trees (black) as well as the axonal arbor (red) for 4 of the 5 cells recovered from whole-cell recordings in slices from CR^Cre^;Ai9 mice. **e**-**g**: examples of the appearance of the cell shown in **d** as seen with the electron microscope. **e** and **f** show axonal boutons belonging to this cell forming axoaxonic synapses onto the central terminals of type I glomeruli (C_I_). In **g**, part of a dendrite of this cell is postsynaptic to another type I central glomerular bouton (arrowhead). Scale bars = 100 μm (**a**-**d**) and 0.5 μm (**e**-**g**).

### Identification of other genetic mouse lines for targeting inhibitory calretinin cells

Since the iCRs account for only ∼15% of the calretinin-expressing cells in the SDH^34, 35^, we searched for other genetically modified mouse lines and crosses that would allow us to target these cells more efficiently.

Many dorsal horn inhibitory interneurons express the neuropeptide nociceptin^18, 19, 41^. We therefore crossed a BAC transgenic line (Pnoc::GFP) in which green fluorescent protein (GFP) is expressed under the transcriptional control of the prepronociceptin (Pnoc) promoter^41, 42^, with a knock-in calretinin-Cre line (CR^Cre^)^43^ and the Ai9 reporter mouse, in which Cre-mediated excision of a STOP cassette leads to permanent expression of the red fluorescent protein tdTomato. We found that 92.6% (89.1-96.3%, n = 3 mice) of GFP-positive cells in laminae I-II in this line were immunoreactive for Pax2, a transcription factor that is restricted to inhibitory interneurons in the dorsal horn^44, 45^, while 76.5% (71.9-80.6%) of cells that were immunoreactive for both Pax2 and calretinin contained GFP (Fig S1). The Pnoc::GFP;CR^Cre^;Ai9 cross therefore allows for visualisation of the majority of iCRs, and was used for some of our electrophysiological experiments (see below).

The transcription factor RAR-related orphan receptor β (Rorb) is expressed by many inhibitory (and some excitatory) neurons in the dorsal horn^46, 47^. During the course of this study we observed that tdTomato-labelled neurons that were captured in a Rorb^CreERT2^;Ai9 cross^46^ included a distinctive population in the SDH. These cells were concentrated in mid-lamina II, extending across the entire width of the dorsal horn (Fig 2). Although there were also many tdTomato-positive cells in deeper laminae, these were separated from those in lamina II, as there were relatively few labelled cells in the inner (ventral) part of lamina II or the dorsal part of lamina III. We found that 93.3% (± 9.0% S.D.; n = 5 mice) of tdTomato-positive cells in lamina II were Pax2-immunoreactive, and that 84.2% (± 7.2%) of these contained calretinin. However, only 29.4% (± 7.2%) of iCRs in lamina II were labelled with tdTomato in the Rorb^CreERT2^;Ai9 mice. Although there were numerous tdTomato-labelled cells in deeper laminae (III-IV) and most of these were inhibitory (Pax2-positive), <10% of these cells were calretinin-immunoreactive. We therefore used the Rorb^CreERT2^;Ai9 cross to target iCRs in some electrophysiological experiments (see below), as well as in anatomical studies to investigate synaptic connectivity.

**Fig 2.**
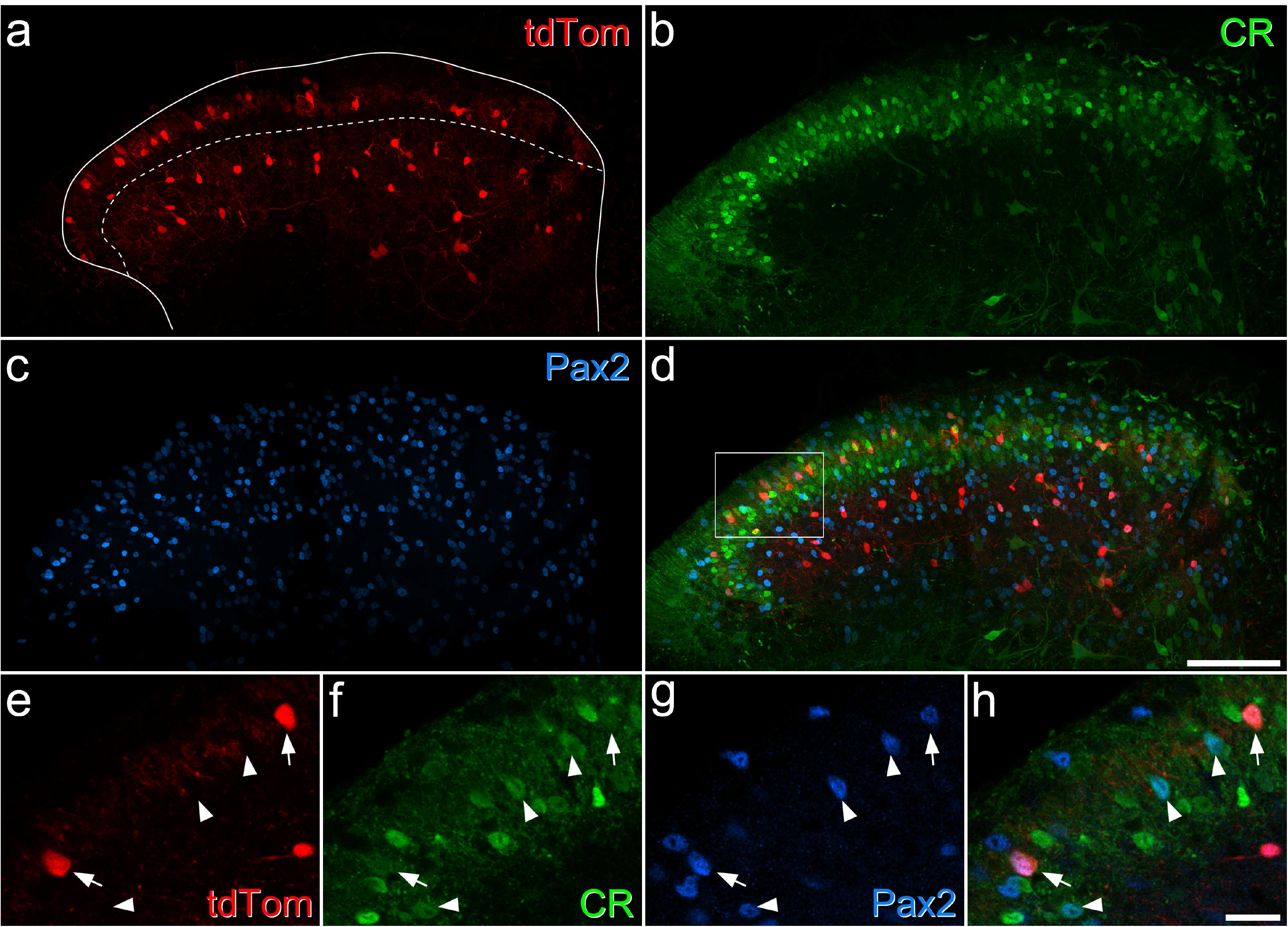
Expression of tdTomato in a Rorb^CreERT2^;Ai9 mouse. **a**-**d**: a transverse section through the lumbar spinal cord from a Rorb^CreERT2^;Ai9 mouse, immunostained to reveal tdTomato (red), calretinin (CR, green) and Pax2 (blue). The solid line shows the outline of the grey matter and the dashed line the approximate position of the lamina II-III border. **a**: there are many tdTomato positive cells in the middle of lamina II, and others in deeper laminae (III-IV), but there are relatively few cells in a narrow band on either side of the border between laminae II and III. **b:** there are many calretinin-immunoreactive cells in laminae I-II, and scattered weakly-labelled cells in the deep dorsal horn. **c**: Pax2 is expressed by all inhibitory interneurons in the dorsal horn, and there are many immunoreactive cells throughout this region. **e**-**h**: a higher magnification view through the superficial laminae (corresponding to the area outlined by the box in **d**). Two tdTomato-positive cells are marked by arrows, and these are both immunoreactive for calretinin and Pax2. Arrowheads mark three other calretinin-positive/Pax2-positive cells that lack tdTomato. Images in **a**-**d** are maximal intensity projections of 24 optical sections at 2 μm z-spacing, while those in **e**-**f** are from a single optical section. Scale bars = 100 μm (**a**-**d**) and 20 μm (**e**-**h**).

Inhibitory calretinin cells belonging to the Gaba9 class of Häring et al^18^ or the DI-1 class of Sathyamurthy et al^19^ express the Tac1 gene, which codes for preprotachykinin A (PPTA), the precursor for substance P. We have previously reported that 23.1% of inhibitory interneurons in SDH express calretinin^34^. In a Tac1^Cre^ mouse line we had found that 11.2% of Tac1-expressing cells in the SDH were inhibitory, while Tac1-expressing cells accounted for 19.6% of all SDH neurons^48^. Since 25.8% of SDH neurons in the mouse are inhibitory^15^, we can estimate that Tac1 cells account for ∼8.5% of SDH inhibitory interneurons. We also found that (as expected from transcriptomic studies^18, 19^) virtually all inhibitory Tac1-expressing cells were calretinin-immunoreactive^35^. Taken together, these results suggest that around 37% of iCRs express Tac1. We therefore used fluorescence *in situ* hybridisation (FISH) to look for *Tac1* mRNA in tdTomato-labelled cells in sections from 3 Rorb^CreERT2^;Ai9 mice. We identified a total of 98 tdTomato-positive cells and found that most (mean 85.5%, range 79.2-86.2%) of these were also positive for *Gad1*, a marker of inhibitory neurons. Among these *tdTomato*+/*Gad1*+ cells, 80.3% (73.7-88.1%) contained *Tac1* mRNA (Fig 3). This suggests that the Rorb^CreERT2^;Ai9 cross captures a subset of iCRs, with preferential labelling of those that coexpress Tac1, and this is consistent with the high level of expression of Rorb in the DI-1 class of Sathyamurthy et al^19^.

**Fig 3.**
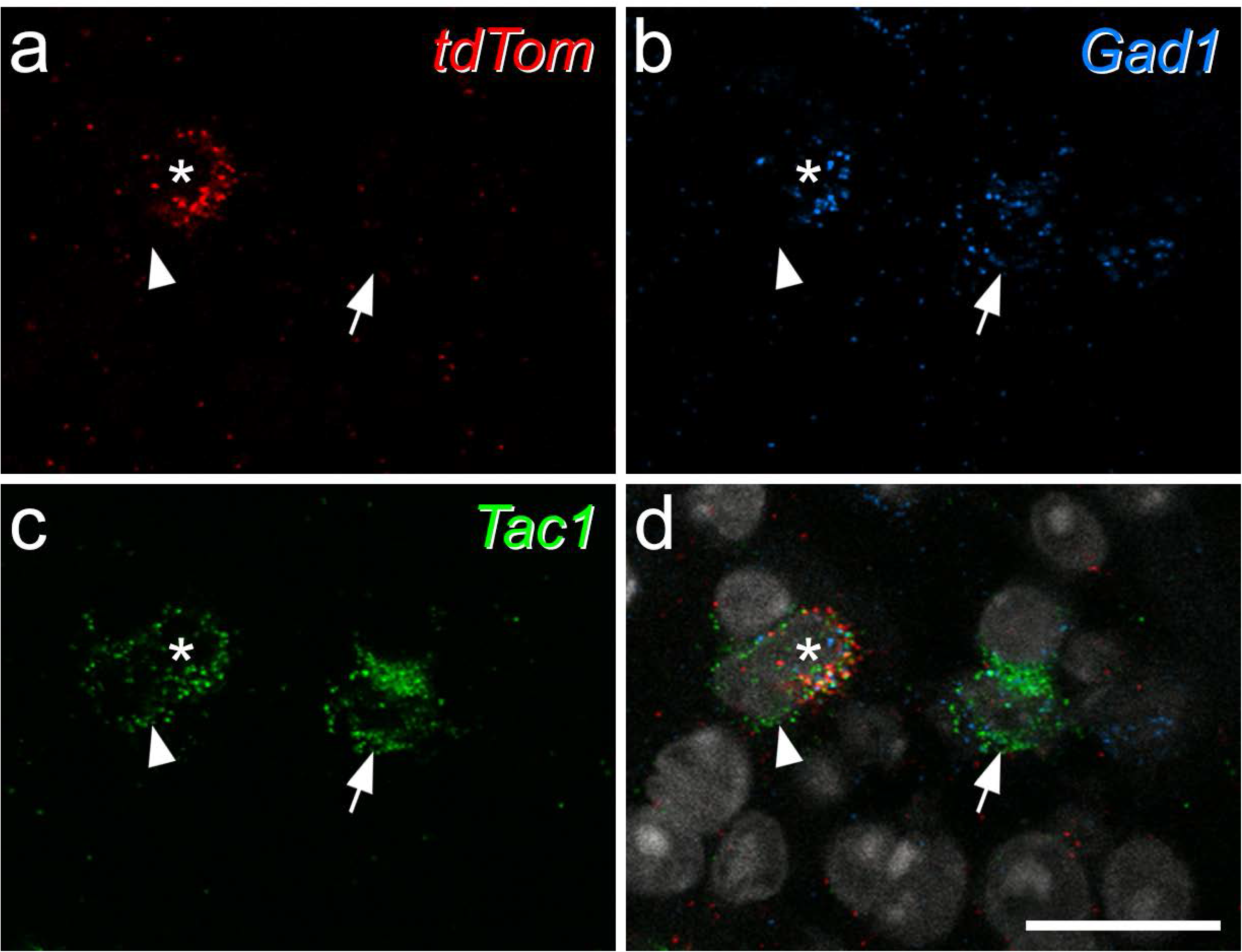
Expression of *Tac1* mRNA in tdTomato-positive neurons in the Rorb^CreERT2^;Ai9 mouse. **a**-**c** show fluorescence *in situ* hybridisation signals for *tdTom* (red), *Gad1*(blue) and *Tac1* (green) mRNAs, while **d** shows a merged image with NucBlue counterstain (grey). The asterisk marks a cell that is positive for all 3 signals, the arrow a *Tac1*+/*Gad1*+ cell that lacks *tdTom*, and the arrowhead an excitatory (*Gad1*-negative) *Tac1*+ cell that also lacks *tdTom*. Images are maximum intensity projections of confocal optical sections (1 μm z-separation) through the full thickness of the section. Scale bar = 20 μm.

Although many iCRs contain *Tac1* mRNA, and we have identified the protein product (PPTA) in some inhibitory interneurons in the SDH, substance P can only be detected at extremely low levels in a few GABAergic boutons in this region^48^. Sathyamurthy et al^19^ reported that the DI-1 cells have a very low level of mRNA for the enzyme peptidylglycine alpha-amidating monooxygenase (PAM), which is required for amidation of substance P. To test whether this could explain the lack of substance P in inhibitory axonal boutons we used FISH to detect transcripts for *Slc32a1* (which encodes the vesicular GABA transporter), *Tac1* and *Pam*. In tissue from 3 wild-type mice we identified a total of 54 inhibitory (*Slc32a1*-positive) and 347 excitatory (*Slc32a1*-negative) *Tac1*-containing cells. The number of transcripts for *Tac1* did not differ between these populations (19.1 ± 12.7 for inhibitory cells, 19.6 ± 13.9 transcripts for excitatory cells; Mann-Whitney U test, p = 0.98). However, transcripts for PAM were significantly less numerous in the inhibitory cells (4.3 ± 3.3, compared to 15.1 ± 11.2 transcripts for excitatory cells; Mann-Whitney U test, p < 0.001) (Fig S2). This suggests that the lack of substance P in axons of these cells results from lack of PAM, which is required for synthesis of the mature peptide, rather than from low expression of Tac1.

### Electrophysiological characterisation of iCRs and their responses to dorsal root stimulation

Based on the findings described above, we recorded from cells in the SDH that co-expressed GFP and tdTomato in tissue from Pnoc::GFP;CR^Cre^;Ai9 mice as well as from tdTomato labelled SDH cells in tissue from Rorb^CreERT2^;Ai9 mice. For some experiments we crossed Rorb^CreERT2^ mice with the Ai32 reporter, in which Cre-mediated excision of a STOP cassette results in expression of a channelrhodopsin-YFP conjugate and recorded from YFP-labelled SDH cells. Whole cell patch-clamp recordings in sagittal slices were made from a total of 200 cells, comprising 82 from Pnoc::GFP;CR^Cre^;Ai9, 107 from Rorb^CreERT2^;Ai9 tissue and 11 from Rorb^CreERT2^;Ai32 mice. No differences were observed between female and male mice, therefore all data presented are a combination of both sexes. Except where stated, no differences were observed between recordings performed in Pnoc::GFP;CR^Cre^;Ai9 and Rorb^CreERT2^ tissue.

In agreement with Smith et al.^34^, we found that the majority of iCRs (which correspond to their “atypical calretinin cells”) exhibited tonic (75/92; 81.5 %) or transient (12/92; 13.0 %) action potential firing patterns in response to depolarising current steps, with a small proportion exhibiting single spike firing (5/92; 5.4 %) (Fig 4a,b). The membrane properties of these cells are detailed in Table 1, and mean values described here. Table 1 also provides a comparison of data obtained from Pnoc::GFP;CR^Cre^;Ai9 tissue with that from Rorb^CreERT2^ tissue. The mean resting membrane potential of iCRs was -58.4 mV, and was significantly more depolarised for cells recorded in Pnoc::GFP;CR^Cre^;Ai9 than Rorb^CreERT2^ tissue (-54.2 vs. -62.0 mV, p<0.0001, unpaired t test). Their input resistance was 959.4 MΩ and their capacitance 10.7 pF. The rheobase current was 17.8 pA, with the following parameters measured from the first action potential to occur at rheobase: the action potential voltage threshold (defined as the point where the rate of rise exceeded 10 mV/ms) was -35.1 mV, the latency between the onset of the depolarising step to the first action potential was 252.3 ms, base width was 1.3 ms, action potential height (measured as the difference between the voltage threshold and the peak of the action potential) 74.9 mV, and after-hyperpolarisation was -30.4 mV. Differences between recordings from Pnoc::GFP;CR^Cre^;Ai9 and Rorb^CreERT2^ tissue were found in action potential height (76.9 vs. 73.2 mV, respectively, p = 0.041, Mann Whitney) and base width (1.2 vs. 1.4 ms, respectively, p = 0.009, Mann Whitney).

**Fig 4.**
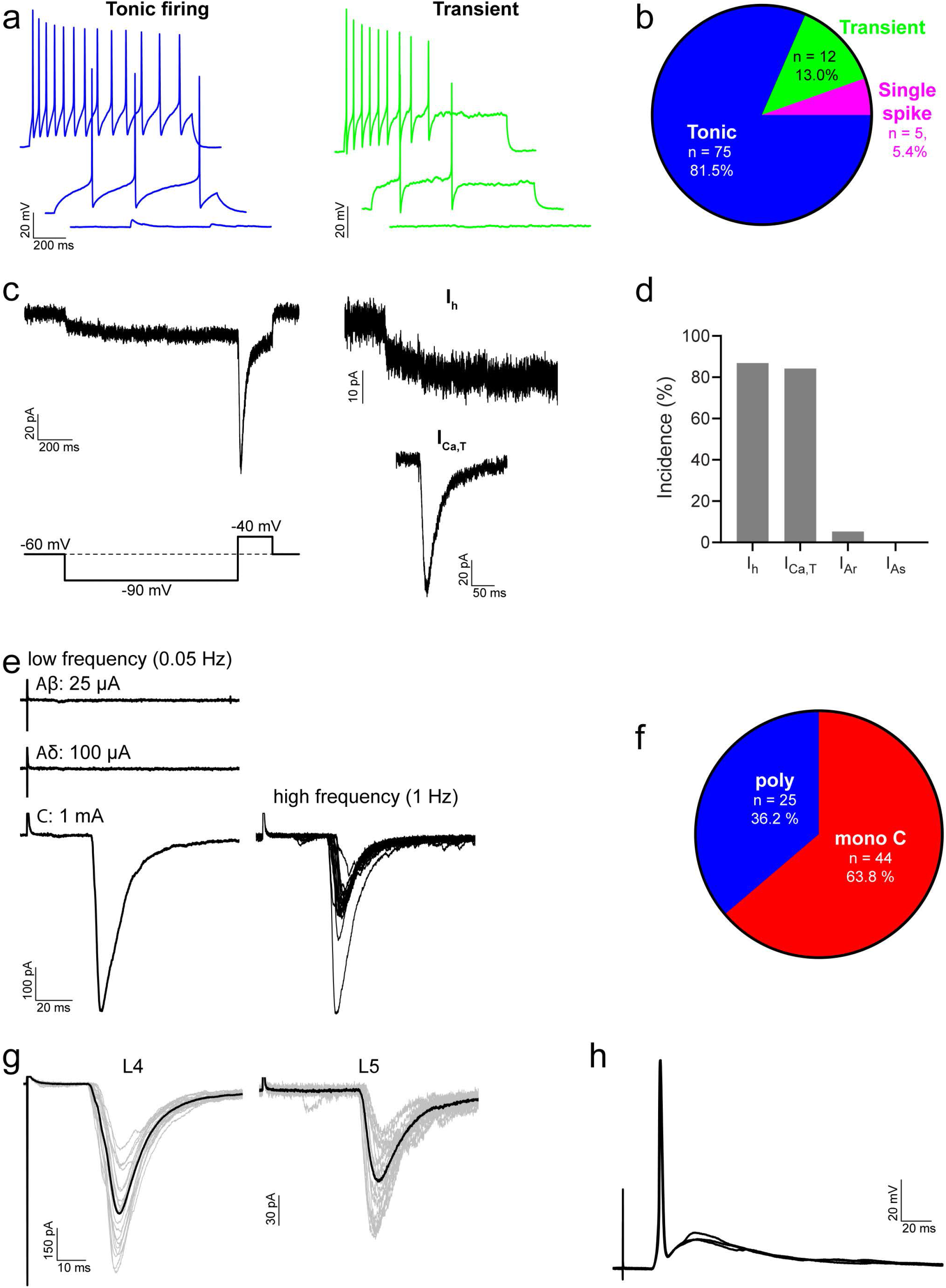
Action potential firing patterns, subthreshold voltage-activated currents and primary afferent input for iCRs. **a** shows examples of action potential firing patterns observed in iCRs in response to 1-second current injections. **b**: most iCRs exhibited tonic firing, with smaller proportions displaying transient or single spike firing. **c**: representative traces showing hyperpolarisation-activated (I_h_) and low-threshold calcium (I_Ca,T_) currents, in response to a voltage step protocol (bottom left). Each trace shows an average of 5 sweeps. **d**: most iCRs displayed I_h_ and/or I_Ca,T_, while A-type potassium currents (I_A_) were rarely observed. **e**: representative traces of monosynaptic C fibre input to iCRs, revealed in response to electrical stimulation of dorsal roots. Low frequency traces are an average of 3 sweeps, high frequency trace shows 20 superimposed sweeps. **f**: the majority of iCRs received primary afferent input that was classified as monosynaptic from C fibres; for the remaining cells the input was classified as polysynaptic only. In six cells we found primary afferent input from both L4 and L5 dorsal roots; an example of a cell with monosynaptic C fibre input from both roots is shown in **g**. Grey traces are 20 individual sweeps, black traces are an average of the grey traces. In most iCRs (8/11) the monosynaptic C fibre input was sufficient to drive action potential firing, and an example is show in **h**. This shows three individual traces superimposed.

**Table 1.**
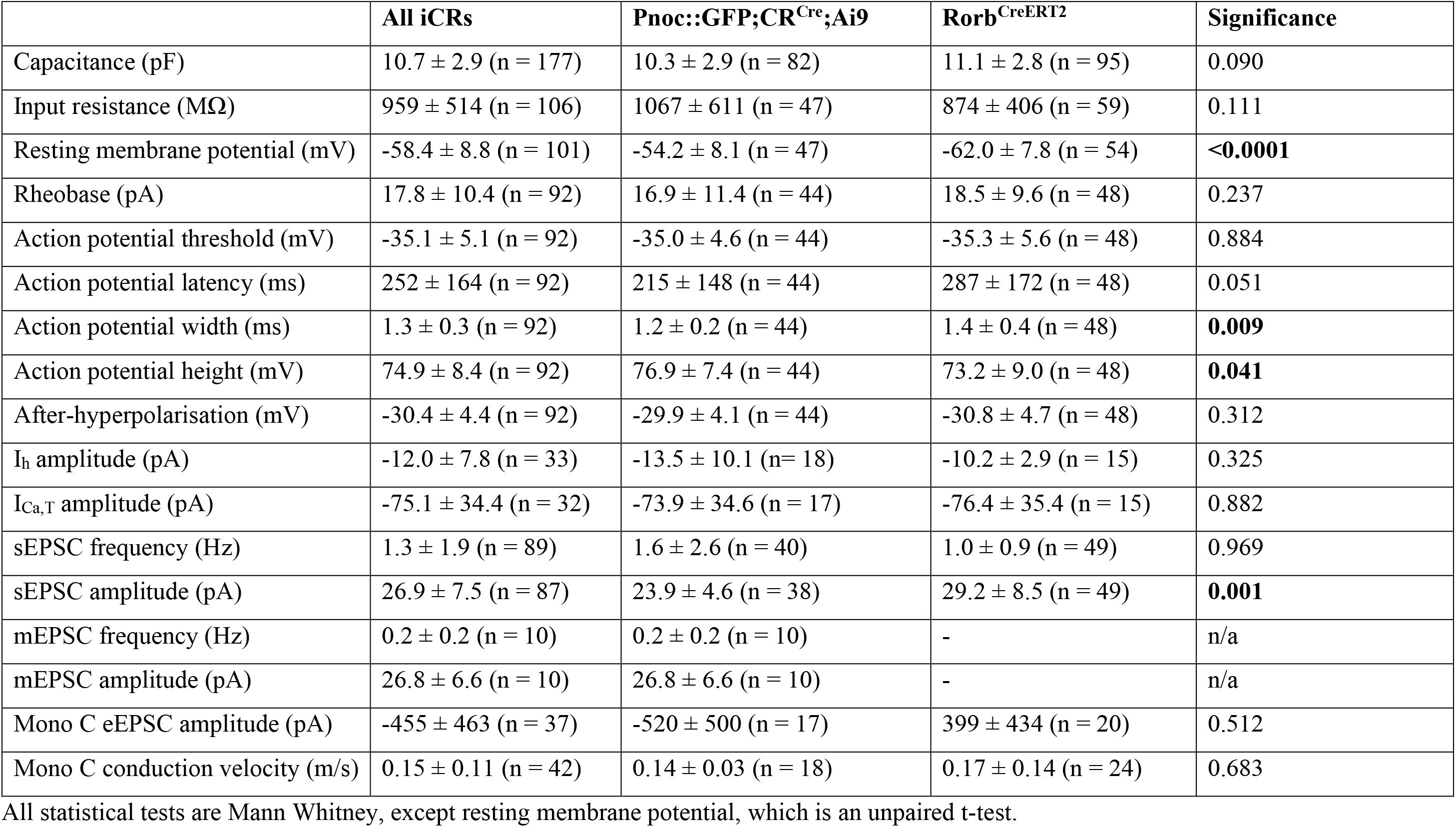
Electrophysiological properties of iCRs and comparison between the properties of Pnoc::GFP;CR^Cre^;Ai9 cells and Rorb^CreERT2^ cells.

Smith et al reported that iCRs exhibit hyperpolarising-activated (I_h_) and “T-type” calcium (I_Ca,T_) subthreshold voltage-activated currents, but do not display A-type potassium currents (I_A_)^34^. Our findings similarly demonstrate that I_h_ (33/38; 86.8 %) and I_Ca,T_ (32/38; 84.2 %) are the dominant subthreshold currents in iCRs, while I_A_ was rarely seen (2/38; 5.3 %) (Fig 4c,d). The amplitude of I_h_ was -12.0 ± 7.8 pA and the peak amplitude of I_Ca,T_ was -75.1 ± 34.4 pA (Table 1).

We investigated excitatory synaptic input to iCRs by recording spontaneous (sEPSC) and miniature (mEPSC) excitatory postsynaptic currents at a holding potential of -70 mV. sEPSC and mEPSC frequencies were 1.3 ± 1.9 and 0.2 ± 0.2 Hz, and the amplitudes were 26.9 ± 7.5 and 26.8 ± 6.6, respectively (Table 1). The sEPSC amplitude from cells in Rorb^CreERT2^ tissue was greater than that of cells in Pnoc::GFP;CR^Cre^;Ai9 tissue (29.2 ± 8.5 vs. 23.9 ± 4.6 pA, p = 0.001, Mann Whitney, Table 1). In some cases we recorded sEPSCs and mEPSCs in the same cell and found that the sEPSC frequency was significantly greater (0.6 ± 0.3 vs. 0.2 ± 0.2 Hz, p = 0.002, Mann Whitney, n=10), suggesting that iCRs receive excitatory synaptic input from other cells in the slice that are spontaneously active.

Primary afferent input to iCRs was investigated in spinal cord slices with dorsal roots attached. Electrical stimulation of the dorsal roots resulted in eEPSCs in 69/89 (77.5 %) cells tested. In the majority of those cells that exhibited eEPSCs (44/69; 63.8 %), this input was classified as monosynaptic C fibre input (Fig 4e,f), with 8 of these cells receiving additional polysynaptic input. The estimated conduction velocity of C fibres providing this monosynaptic input was 0.15 ± 0.11 m/s, and the peak amplitude of the input was -454.5 ± 462.7 pA. In the remaining cells (25/69; 36.2 %) the input that they received was classified as polysynaptic only, the majority of which was revealed during stimulation at C fibre intensity (22 cells), with smaller proportions evoked at Aδ (5 cells) and Aβ (2 cells) intensity. Some cells displayed a mix of different polysynaptic currents. In some recordings we found cells that received input from both L4 and L5 dorsal roots (6 cells, Fig 4g), indicating that iCRs can receive primary afferent input across more than one spinal segment. In current clamp recordings, we found that the activation of monosynaptic C fibre input resulted in action potentials in most cells tested (8/11; 72.7 %, Fig 4h), where typically a single action potential resulted from each stimulus. This demonstrates that the monosynaptic C fibre input to iCRs is sufficiently powerful to drive action potential firing in these neurons.

This monosynaptic C fibre input was further characterised in a subset of cells, to determine whether it originated from primary afferents that express the noxious heat sensing channel, TRPV1. Following bath application of the TRPV1 agonist capsaicin, which blocks action potential firing in TRPV1-expressing afferents^49^, most monosynaptic C fibre input to iCRs was classified as being insensitive to capsaicin (Fig 5a-c), with capsaicin reducing the peak eEPSC amplitude in only 2 out of 14 cells tested (14.3%, Fig 5b,c). This was consistent for recordings made in slices from Rorb^CreERT2^ and Pnoc::GFP;CR^Cre^;Ai9 mice (1 of 6 cells and 1 of 8 cells, respectively), and indicates that the majority of monosynaptic C fibre input to iCRs arises from primary afferents that lack TRPV1. Of the two inputs that were classified as ‘sensitive’, capsaicin reduced the peak eEPSC amplitude from -245.4 to -107.1 and -511.1 to -217.6 pA, while in capsaicin ‘insensitive’ cells the corresponding values were -551.7 ± 400.6 for baseline and -493.2 ± 341.3 pA for capsaicin application. Surprisingly, when we recorded mEPSCs (Rorb^CreERT2^ tissue only), we found that capsaicin resulted in a significant leftwards shift in the cumulative distribution of inter-event intervals, signifying an increase in mEPSC frequency, in almost all cells (7/8; 87.5%, Fig 5d-f). In those cells that responded to capsaicin, the mEPSC frequency was increased from 0.2 ± 0.1 to 5.9 ± 5.1 Hz (p = 0.016, Wilcoxon matched-pairs signed rank test), while in the single cell that did not respond the baseline and capsaicin frequencies were 0.04 and 0.03 Hz, respectively (Fig 5g). The percentage of iCRs that responded to capsaicin here was greater than that previously reported by Smith et al.^34^ (4/7; 57.1 %), as was the magnitude of the increase in mEPSC frequency.

**Fig 5.**
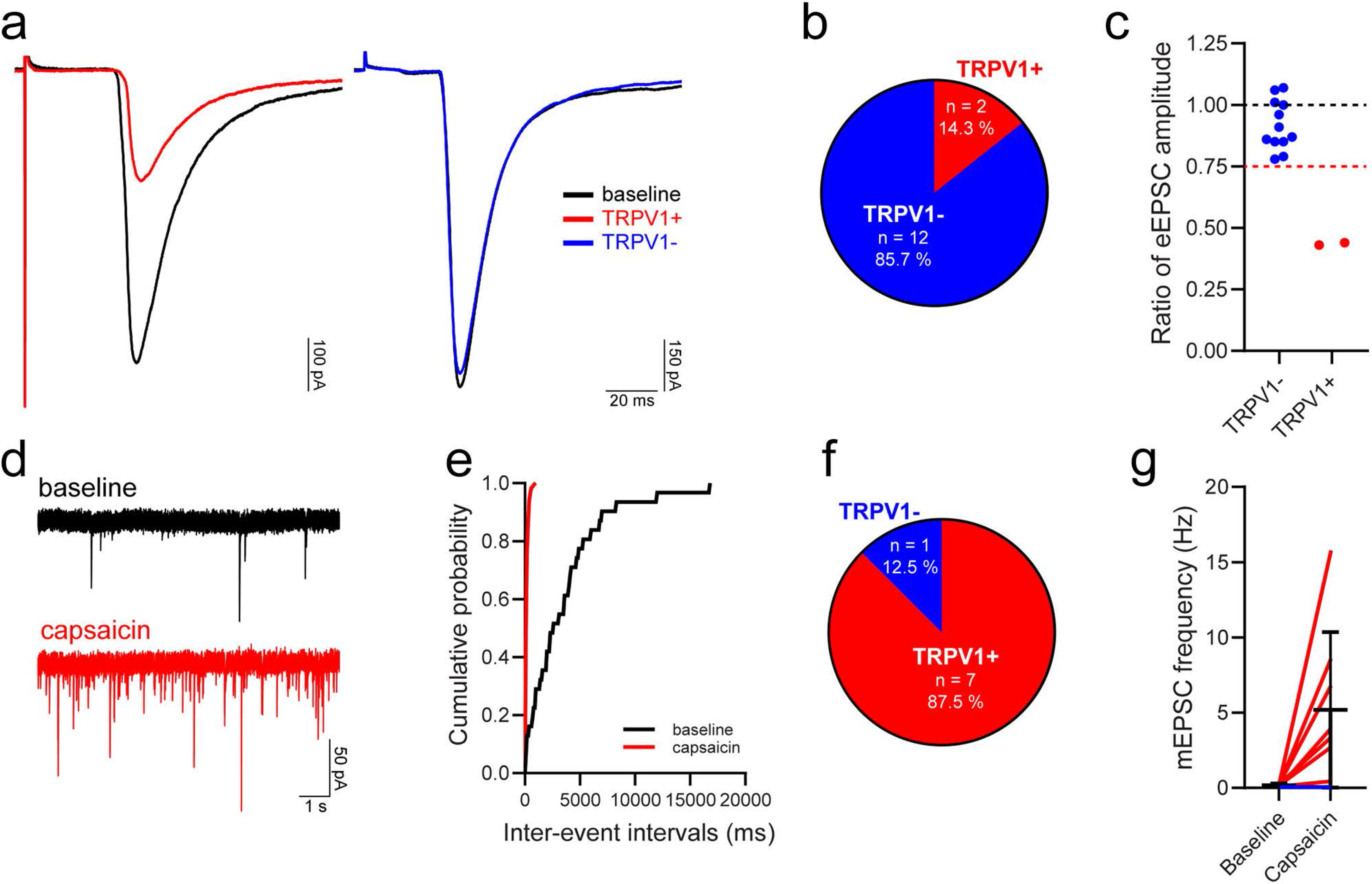
TRPV1 expression by primary afferents that provide input to iCRs. **a**: representative traces showing monosynaptic C fibre input to iCRs before (baseline, black traces) and during application of the TRPV1 agonist capsaicin in two individual cells classified as having TRPV1-sensitive (TRPV1+, red trace) and TRPV1-insensitive (TRPV1-, blue trace) monosynaptic C fibre input. Traces are an average of 15 sweeps, comprising the last 5 minutes of baseline recordings (baseline) or the final 5 minutes of capsaicin application (TRPV1+/TRPV1-). **b,c**: most iCRs received monosynaptic C fibre input that was classified as TRPV1-insensitive. The dashed red line in **c** denotes a value of 0.75, the threshold for defining whether a monosynaptic C fibre input was classified as TRPV1-insensitive (>0.75) or TRPV1-sensitive (≤0.75). **d**: representative mEPSC traces recorded during baseline and during the application of capsaicin. **e**: example of a cumulative probability plot that demonstrates a significant leftward shift in the distribution of mEPSC inter-event intervals in response to the application of capsaicin (P<0.00001, Kolmogorov– Smirnov 2-sample test, taken from the same cell as **d**). **f**: a significant leftward shift in inter-event intervals, signifying an increase in mEPSC frequency, was observed in 7 out of 8 cells tested, with those cells being classified as receiving TRPV1+ primary afferent input. The effect of capsaicin on mEPSC frequency in cells receiving TRPV1+ input (red lines) and the single cell with input that was defined as TRPV1-(blue line) is shown in **g**.

Smith et al.^36^ reported that all iCRs tested demonstrated robust outward currents in response to the MOR agonists, enkaphalin and DAMGO, as well as to the DOR agonist, DADLE. In agreement, when we applied DAMGO to iCRs (Rorb^CreERT2^ tissue only), this resulted in a clear, slow outward current in all cells (7/7; Fig 6a, upper trace). These currents were often large (up to ∼100pA), with a mean amplitude of 50.3 ± 36.7 pA. In contrast, few cells responded to the DOR agonist, [D-Ala2]-Deltophin II, with only 2/8 (25.0 %) cells exhibiting an outward current (Fig 6a, middle trace). We also tested responses to the KOR agonist, U69593, which resulted in an outward current in just over a third of cells (3/8; 37.5%, Fig 6a, lower trace). The outward currents observed in response to Deltorphin II and U69593 were typically smaller (9.9 ± 0.3 pA and 16.0 ± 9.4 pA, respectively) than the DAMGO-mediated currents. These findings demonstrate that iCRs are strongly inhibited by MOR agonists, and that some are inhibited to a lesser extent by DOR and KOR agonists. The findings with DAMGO are consistent with transcriptomic data, which shows expression of *Oprm1* (the gene encoding MOR) among at least some cells in the iCR clusters^18, 19^. This seems paradoxical, since opioids, which are potent analgesics, will therefore suppress activity in inhibitory interneurons. However, it is likely that this effect is outweighed by other actions of the opioids on both primary afferents and other CNS neurons.

**Fig 6.**
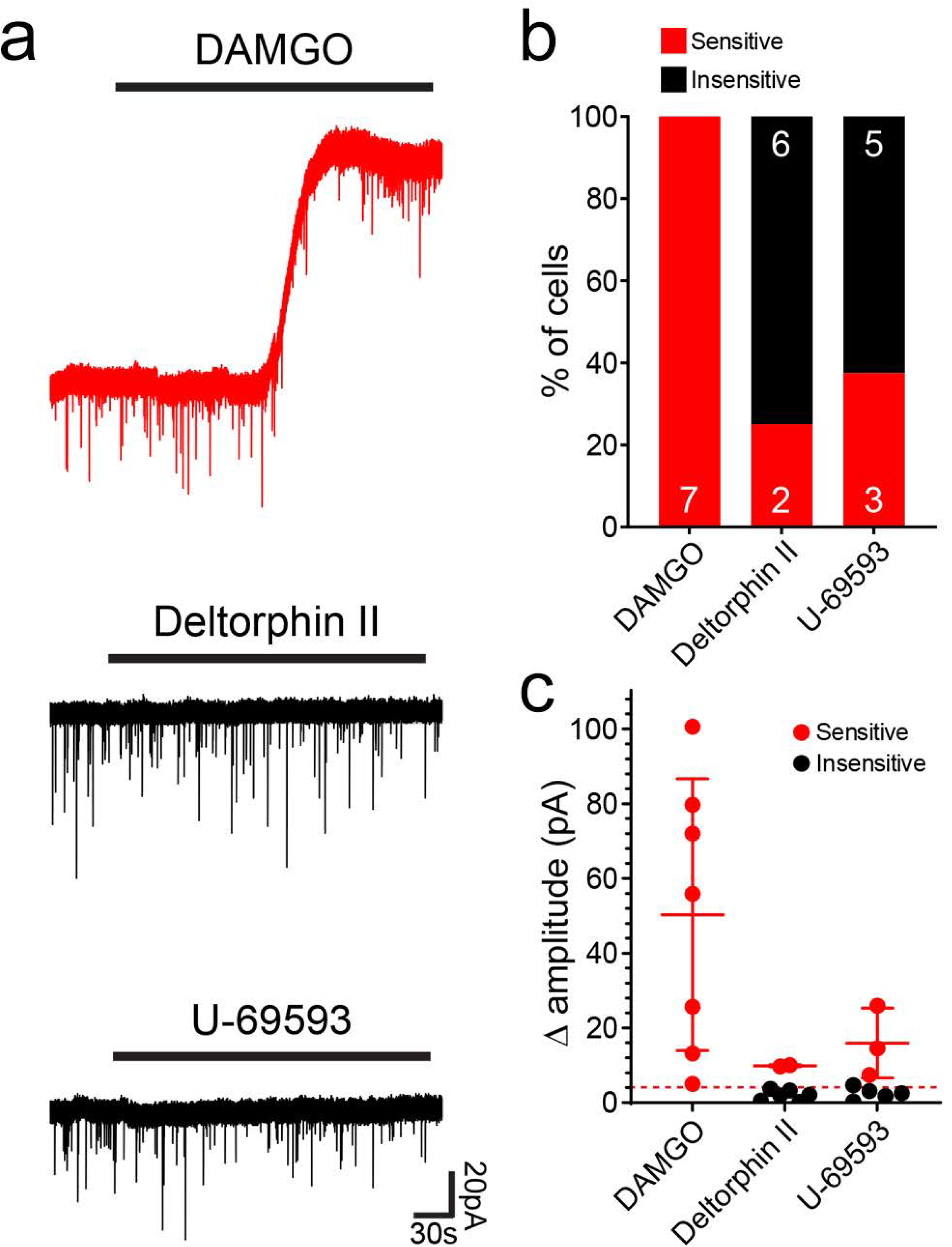
Responses of iCRs to opioids tested in Rorb^CreERT2^;Ai9 mice. **a**: examples of responses to the μ-opioid agonist DAMGO, the δ-opioid agonist deltorphin II and the κ-opioid agonist U-69593. The cell tested with DAMGO showed an outward current (top trace), while those tested with Deltorphin II (middle trace) and U-69593 (bottom trace) did not respond. **b**: the proportions of cells that were sensitive or insensitive to each agonist. **c**: the amplitudes of responses to the agonists for each cell tested, with mean and SD of the cells sensitive to each agonist. The dashed line represents a value of 5 pA, which was taken as the threshold for defining a responsive cell.

### Synaptic connectivity of Rorb cells in lamina II

Since the Rorb^CreERT2^ mouse was found to capture a subset of iCRs, we performed multiple-labelling immunohistochemistry and confocal microscopy on tissue from crosses involving this line to investigate synaptic inputs to, and outputs from, these cells.

We used the MrgD^ChR2-YFP^ mouse line, in which channelrhodopsin fused to yellow fluorescent protein (YFP) is knocked into the MrgD locus^50^, to identify non-peptidergic nociceptors belonging to the NP1 population^9^. This was crossed with the Rorb^CreERT2^ and Ai9 lines to generate Rorb^CreERT2^;Ai9;MrgD^ChR2-YFP^ mice, in which Rorb cells and MrgD afferents were labelled with tdTomato and YFP, respectively. The lectin IB4 binds mainly to non-peptidergic nociceptors, including those belonging to NP1 and NP2 classes, but not to NP3 afferents, C-low threshold mechanoreceptors (C-LTMRs) or to many peptidergic nociceptors^2, 51–56^. We therefore used IB4 binding and also identified IB4-positive YFP-negative profiles, which are likely to correspond to NP2 central terminals.

Transverse sections from the L4 segments of these mice were immunoreacted with antibodies to reveal tdTomato, YFP and the postsynaptic density marker Homer^46, 57^, and incubated in biotinylated IB4, which was revealed with avidin conjugated to Pacific Blue. We identified an average of 297 YFP-labelled boutons (228-354; n = 3 mice) and found that 74.6% (68.0-87.3%) of these boutons were in contact with at least one Homer punctum that was associated with a tdTomato-labelled profile, while 40.6% (33.8-52.5%) contacted two or more puncta associated with tdTomato-labelled profiles (Fig 7a-d). For the IB4+/YFP-afferents, we found a mean of 28 (21-36; n = 3 mice) boutons, of which 61.2% (30.6-81.5%) were in contact with at least one Homer punctum associated with a tdTomato profile and 35.3% (13.9-47.6%) contacted more than one Homer punctum associated with a tdTomato profile (Fig 7e-h). To identify NP3 afferents we immunostained for somatostatin (SST) and prostatic acid phosphatase (PAP)^58^ on transverse sections of the L4 segment from Rorb^CreERT2^;Ai9 mice. We identified an average of 31 (26-39) SST+/PAP+ boutons (n = 3 mice), of which 71.4% (59.0-82.1%) were in contact with at least one Homer punctum associated with a tdTomato profile and 25.8% (18.0-30.8%) contacted more than one Homer punctum associated with a tdTomato profile (Fig 7i-l). Although there are many tdTomato-expressing cells in the deep dorsal horn of Rorb^CreERT2^;Ai9 mice, there was always a clear gap in the labelling between that in the SDH and that in deeper laminae (Fig 2a), suggesting that the deep cells did not have dendrites extending dorsally into the SDH. It is therefore likely that tdTomato-labelled profiles seen in lamina II originated from cells in the superficial region. These findings therefore suggest that iCRs labelled in the Rorb^CreERT2^;Ai9 mouse receive direct synaptic input from all 3 classes of non-peptidergic nociceptors, and that these cells represent a significant target for NP1-3 afferents.

**Fig 7.**
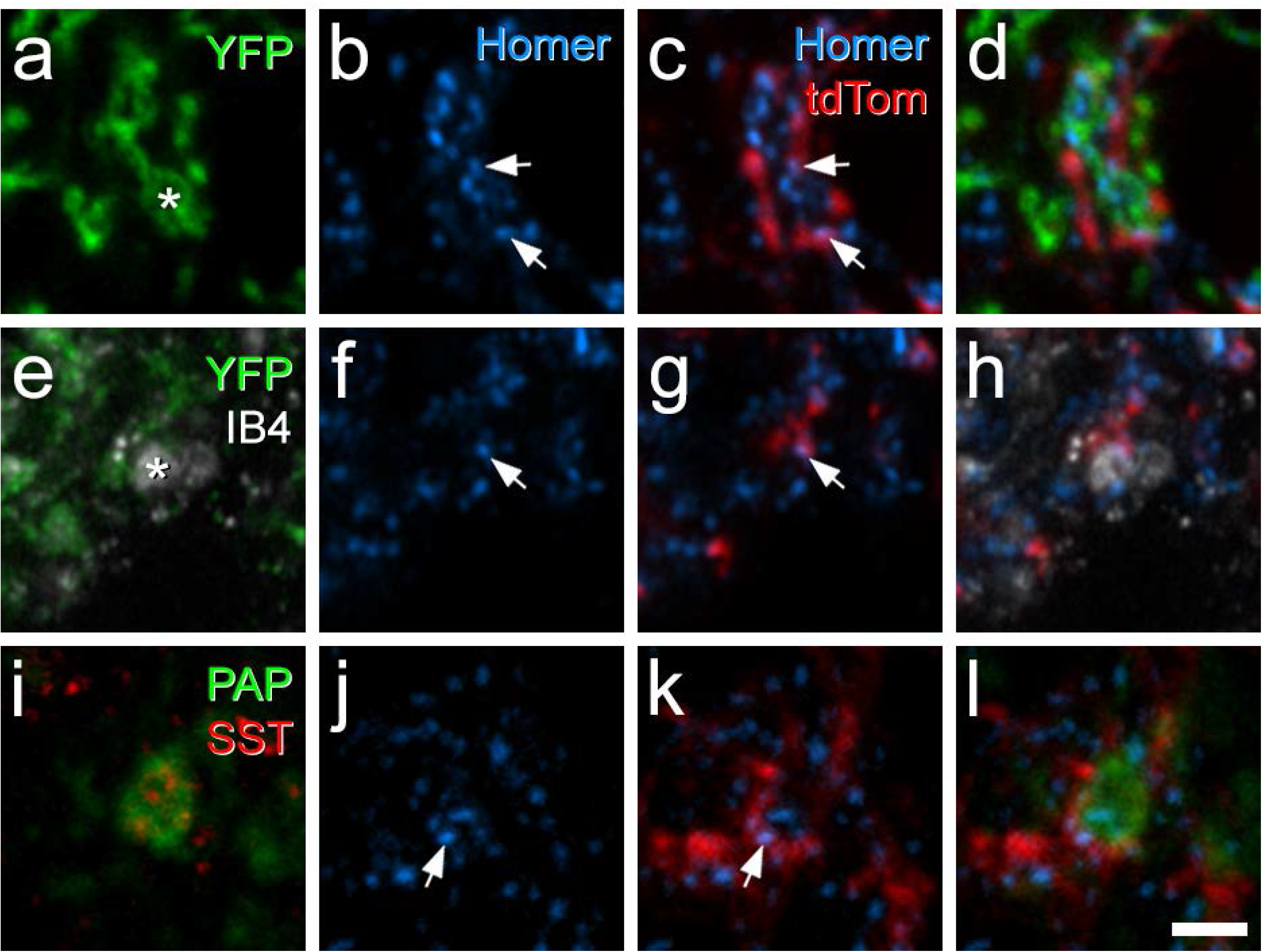
Confocal images showing synaptic input to Rorb cells from non-peptidergic nociceptors. **a**-**d**: immunostaining for YFP (green), Homer (blue) and tdTomato (red) in lamina II in a section from a Rorb^CreERT2^;Ai9;MrgD^ChR2-YFP^ mouse. A YFP-positive central terminal (marked with asterisk in **a**) is surrounded by several Homer puncta. Two of these (marked with arrows) are in tdTomato-positive profiles. **e**-**h**: a region from lamina II in another Rorb^CreERT2^;Ai9;MrgD^ChR2-YFP^ mouse, scanned to reveal YFP (green), IB4 binding (grey), Homer (blue) and tdTomato (red). A central terminal that binds IB4 but lacks YFP is indicated with an asterisk in **e**. This is adjacent to two Homer puncta, one of which (marked with arrows in **f** and **g**) is in a tdTomato-labelled profile. **i**-**l**: immunostaining for prostatic acid phosphatase (PAP, green) and somatostatin (SST, red) reveals the central terminal of a SST-expressing (NP3) afferent. This is in contact with a Homer punctum (arrows in **j** and **k**) that is located in a tdTomato-labelled profile. Images in **a**-**d** and **i**-**k** are from single confocal optical sections, while those in **e**-**h** are from a projection of 3 optical sections at 0.3 μm z-separation. Scale bar = 2 μm.

To identify vesicle-containing profiles belonging to the Rorb cells, we crossed the Rorb^CreERT2^ mouse with the Ai34 reporter line, in which Cre-dependent excision of a STOP cassette results in expression of a tdTomato-synaptophysin fusion protein that is targeted to synaptic vesicles. Since synaptic vesicles are present in dendrites of lamina II islet cells^59, 60^ tdTomato labelling in this cross will capture both axonal boutons and vesicle-containing dendrites (VCDs). We then generated Rorb^CreERT2^;Ai34;MrgD^ChR2-YFP^ mice to investigate the relationship between these vesicle-containing structures and YFP-labelled MrgD afferents. In these mice, tdTomato-labelled profiles formed a dense plexus that occupied lamina II (Fig S3a), and was co-extensive with the band of MrgD afferents (Fig S3b-d). There were also many tdTomato profiles in laminae III and IV, although these were generally separated from the superficial plexus by a region with little tdTomato expression. Based on comparison with the distribution of tdTomato-labelled cell bodies in the Rorb^CreERT2^;Ai9 mice (Fig 2a) and the location of axonal arbors of iCRs (Fig 1a-d), it is likely that this superficial plexus was derived from the Rorb cells in lamina II, while labelled profiles in laminae III-IV originated from Rorb cells in these laminae.

We identified a total of 463 (215, 248, n = 2 mice) MrgD-labelled boutons in the SDH and found that 68.8% of them (61.0%, 76.6%;) received at least one contact from a tdTomato-labelled profile (Fig 8a), while 27.5% (24.7%, 30.2%) were contacted by more than one of these profiles. We analysed 60 (29, 31) IB4-labelled YFP-negative boutons in this tissue (Fig 8b) and found that 78.5% (82.8%, 74.2%) had at least one contact from a tdTomato profile, with 23.4% (24.1%, 22.6%) receiving more than one contact. We also identified 90 (54, 36) somatostatin afferent boutons, identified by co-localisation of SST and PAP immunoreactivity (Fig 8c). Within this population, 35.7% (40.7%, 30.6%) received at least one contact from a tdTomato profile and 8.8% (9.3%, 8.3%) received more than one contact.

**Fig 8.**
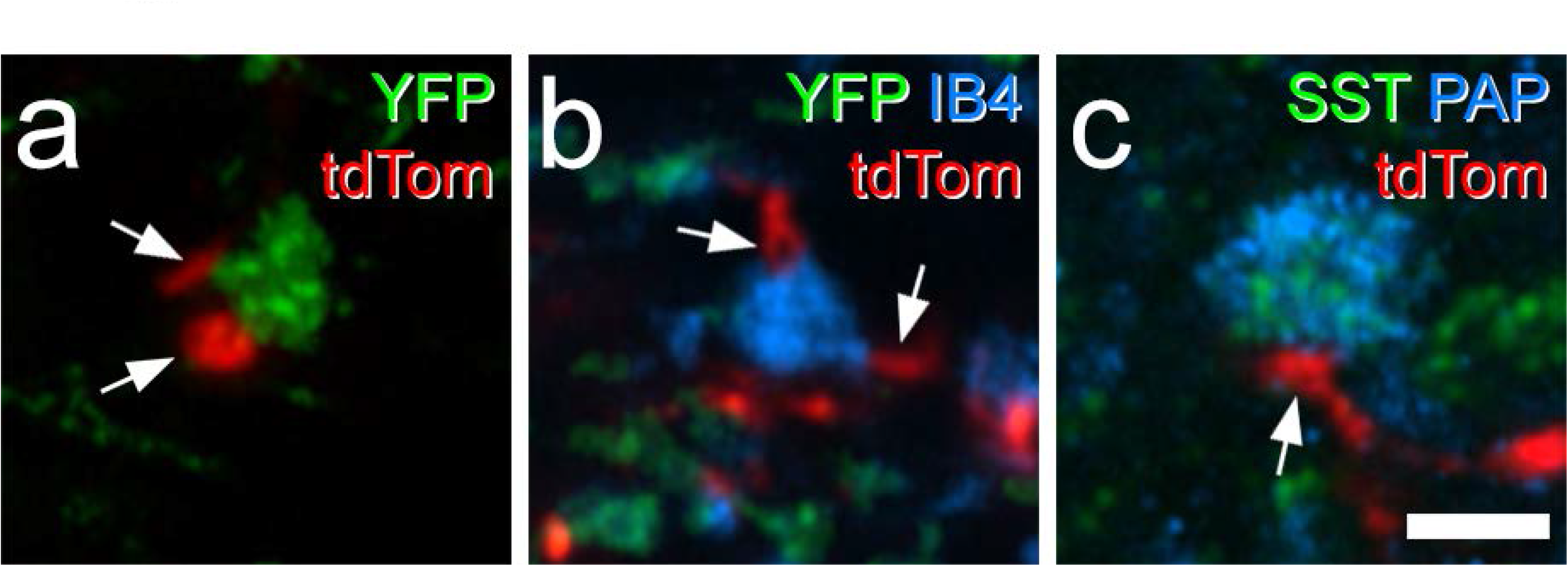
Contacts between Rorb axons and central terminals of non-peptidergic nociceptors in lamina II seen in Rorb^CreERT2^;Ai34;MrgD^ChR2-YFP^ mice. **a**: immunostaining for YFP (green) and tdTomato (tdTom, red) reveals a YFP-positive (MrgD, NP1) central terminal that receives contacts from two tdTomato-labelled profiles (arrows). **b**:in this case, IB4 binding (blue) is seen in a central terminal that lacks YFP (green), and probably belongs to a NP2 afferent. This receives two contacts from tdTomato-labelled profiles (arrows). **c**: immunostaining for somatostatin (SST, green) and prostatic acid phosphatase (PAP, blue) is seen in the central terminal of a SST-expressing (NP3) primary afferent. This is in contact with a tdTom-labelled profile (arrow). All images are projections of 3 optical sections at 0.3 μm z-spacing. Scale bar = 2 μm.

We also used transverse sections from the Rorb^CreERT2^;Ai34;MrgD^ChR2-YFP^ mice to determine the proportion of Rorb-derived vesicle-containing profiles that contacted a MrgD central terminal. For this analysis we used the overlap of IB4 binding and YFP immunoreactivity to define these terminals, because YFP was also present on intervaricose portions of the MrgD axons. We identified 100 tdTomato labelled profiles in the SDH of each of these mice and found that in both cases 70% (n = 2 mice) were in contact with a MrgD central terminal, suggesting that these primary afferent terminals are a major synaptic target of the Rorb cells in lamina II (Fig S3b-d).

GABAergic boutons and VCDs can be presynaptic to central (primary afferent) terminals in glomeruli, forming axoaxonic and dendroaxonic synapses, respectively. They can also form axodendritic and dendrodendritic synapses on dendrites in the glomerulus, and these often contribute to synaptic triads^4, 61^. Inhibitory synapses on dendrites can be identified with confocal microscopy by the presence of the receptor-anchoring protein gephyrin^62, 63^, whereas axoaxonic and dendroaxonic synapses are not associated with gephyrin^64^. We therefore examined sections from 2 Rorb^CreERT2^;Ai34 mice that had been reacted to reveal tdTomato, the vesicular GABA transporter (VGAT, a marker for GABAergic/glycinergic boutons) and gephyrin. We found that on average 91.9% (89.5%, 94.3%) of tdTomato-labelled profiles contained detectable VGAT, and that 73.2% (70.2%, 76.1%) of these were apposed to gephyrin puncta (Fig S3e-h). Taken together, our findings therefore suggest that in addition to forming axoaxonic and dendroaxonic synapses on non-peptidergic nociceptor terminals, the majority of Rorb-derived axons and VCDs also form axodendritic/dendrodendritic synapses, many of which are likely to be components of synaptic triads.

### Intersectional targeting of iCRs

To target the iCRs directly, we crossed CR^Cre^ mice with the VGAT^Flp^ line^65^, in which Flp recombinase is expressed by all inhibitory neurons. We then made intraspinal injections of an AAV that coded for GFP in the presence of both Cre and Flp (AAV.C^on^/F^on^.GFP) into CR^Cre^;VGAT^Flp^ mice. This resulted in expression of GFP that was largely restricted to lamina II, with a few labelled cells in deeper laminae (Fig 9a, b). In parallel experiments, we also injected AAV.C^on^/F^on^.GFP into Tac1^Cre^;VGAT^Flp^ mice, in order to restrict labelling to the Gaba9 population of Häring et al^18^ and the DI-1 class of Sathyamurthy et al^19^. This resulted in a very similar expression pattern, and again GFP-labelled cells and their processes were largely limited to lamina II (Fig S4).

**Fig 9.**
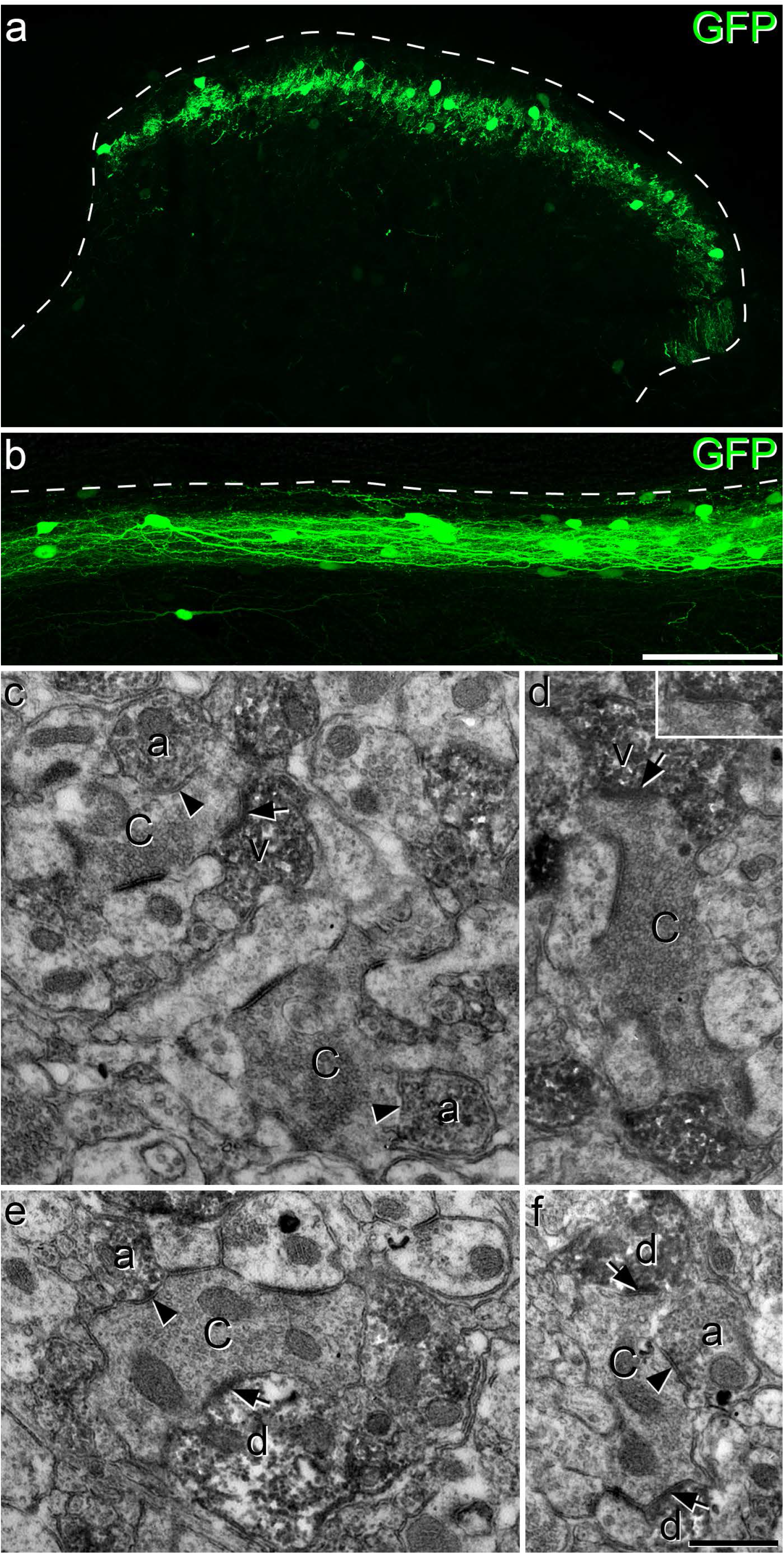
Confocal and electron microscopy of iCRs identified in a CR^Cre^;VGAT^Flp^ mouse injected with AAV.C^on^/F^on^.GFP. **a** and **b:** transverse and sagittal sections demonstrate that GFP labelling is largely restricted to lamina II. In each case the dashed line represents the dorsal border of the grey matter. **c**-**f**: EM images from lamina II obtained following an immuno-peroxidase/DAB reaction to reveal GFP. **c**: two glomerular central terminals (C) are contacted by several DAB-labelled profiles. Two of these (a) contain numerous synaptic vesicles and form axoaxonic synapses (marked by arrowheads) onto the central terminals. One of the DAB-labelled profiles (v) is a vesicle-containing dendrite (VCD) that receives a synapse (arrow) from one of the central terminals. **d**: a glomerular central terminal (C) is in contact with 2 DAB-labelled profiles. One of these (v) can be identified as a VCD, as it contains vesicles and receives an asymmetrical synapse (arrow) from the central terminal. The inset shows the synaptic cleft, as seen following tilting of the specimen. Note that the central terminals in both **c** and **d** resemble those in type I glomeruli, as defined by Ribeiro-da-Silva and Coimbra^4^. **e**,**f** show association between DAB-labelled profiles and glomerular central axons that do not show typical features of those in type I glomeruli. In each case the central terminal (C) receives an axoaxonic synapse (arrowhead) from a DAB-labelled profile identified as an axon terminal (a) and is presynaptic to at least one DAB-labelled dendrite (d) at an axodendritic synapse (arrows). Images in **a** and **b** are projections of 9 and 27 confocal optical sections at 1 μm z-spacing. Scale bars = 100 μm (**a**,**b**) and 0.5 μm (**c**-**f**).

We used the EM to examine sections from CR^Cre^;VGAT^Flp^ mice that had been injected with AAV.C^on^/F^on^.GFP, and then immunoreacted to reveal GFP with an immunoperoxidase method. As expected, we found numerous peroxidase-labelled profiles in lamina II. Many of these profiles were in contact with unlabelled axon terminals that formed multiple axodendritic synapses, corresponding to the central terminals of synaptic glomeruli^4, 61, 66, 67^. Some of these central terminals had a relatively dark appearance, highly indented contours, few mitochondria, and vesicles of irregular size (Fig 9c,d) and were similar to those identified as belonging to the type I glomeruli described by Ribeiro-da-Silva and colleagues^4, 61^. However, many of the central terminals differed from this pattern, for example in terms of electron density, consistency of vesicle size and number of mitochondria (Fig 9e,f; Fig 10). Often more than one peroxidase-labelled profile was in contact with a single glomerular central axon (Fig 9c-f; Fig 10).

**Fig 10.**
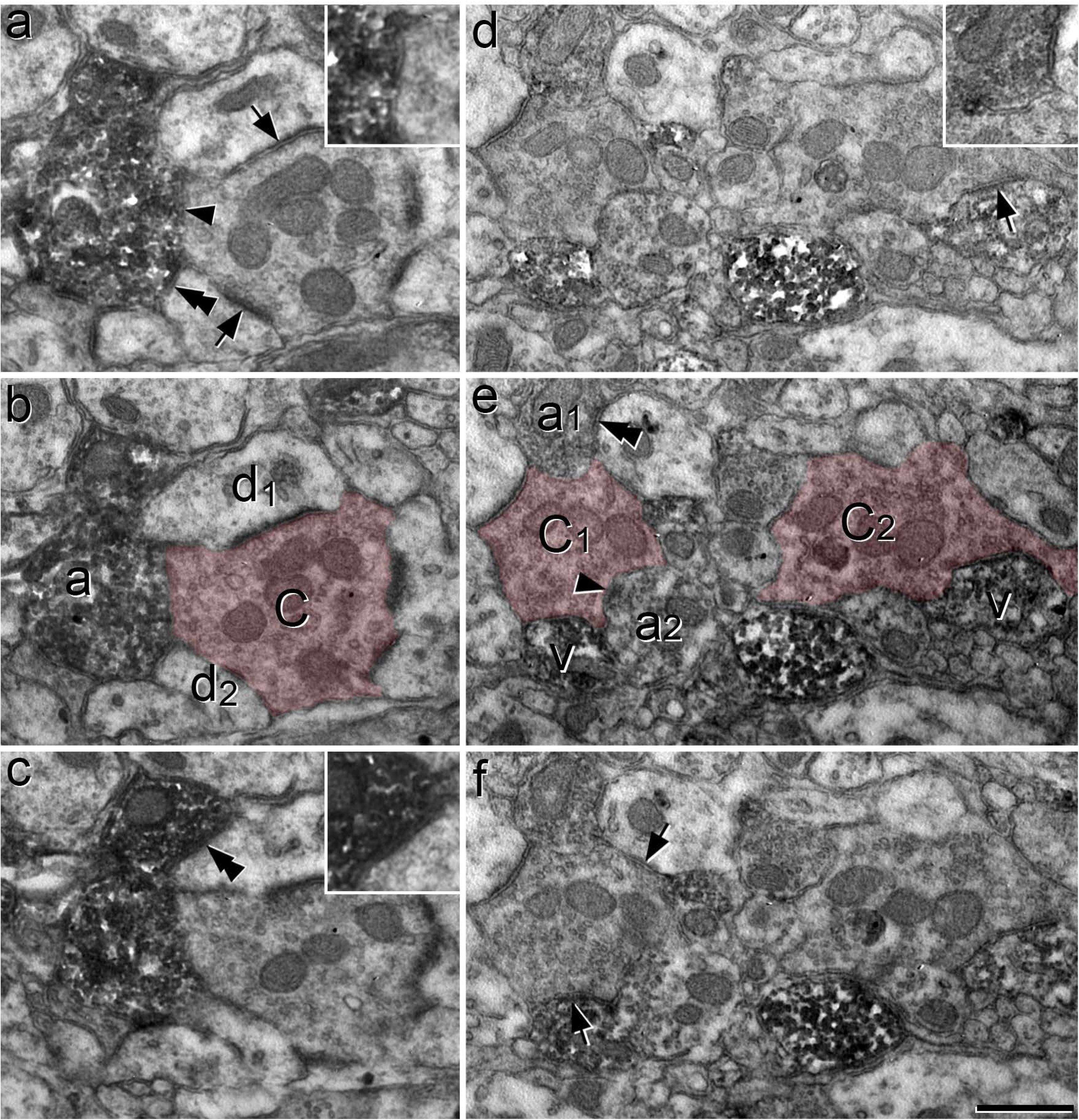
Synaptic triads involving iCRs, as seen in serial ultrathin sections from a CR^Cre^;VGAT^Flp^ mouse that had received intraspinal injection of AAV.C^on^/F^on^.GFP. **a**-**c**: serial ultrathin sections show a glomerular central terminal (C) that receives an axoaxonic synapse (arrowhead in **a**) from a DAB-labelled axonal bouton (a). This synapse is slightly oblique to the plane of section and is seen more clearly following tilting of the specimen (inset in **a**). The DAB-labelled axon is also presynaptic to two dendrites (d_1_, d_2_) at synapses marked by double arrowheads. The synapse on d_1_ is shown in an enlarged view in the inset in **c**. The central terminal is presynaptic to both d_1_ and d_2_ at synapses that are marked with arrows in **a**. These therefore represent triadic arrangements, in which an inhibitory axon forms axoaxonic synapses onto the central terminal and axodendritic synapses onto dendrites that are also postsynaptic to the central terminal. **d-f**: serial ultrathin sections through another region in lamina II include the central terminals of two nearby synaptic glomeruli (C_1_, C_2_), which are associated with several DAB-labelled structures. The terminal marked C_1_ is presynaptic to a labelled VCD (v) at an axodendritic synapse (arrow in **f**), and receives axoaxonic synapses from two labelled axonal boutons (a_1_, a_2_). One of these synapses (from a_2_) is indicated with an arrowhead in **c**, while the other (from a_1_) can be seen in the inset in **d**. The bouton a_1_ also forms a synapse onto an unlabelled dendrite (double arrowhead in **e**) and this dendrite receives a synapse from the C_1_ terminal (arrow in **f**), completing a synaptic triad. The other central terminal (C_2_) is presynaptic to a labelled VCD (v) at an axodendritic synapse (arrow in **d**). The central terminals are indicated with pink shading in **b** and **e**. Scale bar = 0.5 μm.

Many of the peroxidase-labelled profiles in these glomeruli contained synaptic vesicles. However, dendrites in synaptic glomeruli often contain vesicles^4, 61, 67^, and it was therefore sometimes difficult to determine in single ultrathin sections whether the peroxidase-labelled profiles were axonal boutons or VCDs. The main distinction between these structures is that the VCDs can be postsynaptic to the glomerular central terminal, whereas the axonal boutons are not^4, 61^. Based on this criterion, we could identify some of these labelled structures as VCDs (structures labelled “v” in Figs 9c,d and 10e). In many other cases the labelled vesicle-containing profile was presynaptic, but not postsynaptic to the unlabelled central terminal, and was therefore identified as an axonal bouton (structures labelled “a” in Fig 9c,e,f and 10b,e). In many cases these boutons were also presynaptic to dendrites in the glomeruli that received synaptic input from the central terminal, forming synaptic triads (Fig 10). Overall, these observations indicate that many profiles belonging to iCRs form components of synaptic glomeruli. These include axonal boutons that form axoaxonic synapses, often as part of a synaptic triad, as well as VCDs that are postsynaptic to the central bouton.

### Electrophysiological evidence for presynaptic inhibition

GABA released at axoaxonic synapses causes depolarisation of the postsynaptic primary afferent central terminal (primary afferent depolarisation, PAD)^7^. Under certain experimental conditions, notably when recording temperature is reduced below physiological levels, this can lead to the release of glutamate and the appearance of EPSCs in dorsal horn neurons innervated by the affected primary afferent^23, 25, 68^. We therefore carried out optogenetic experiments in CR^Cre^;Ai32 mice, in which a channelrhodopsin-YFP fusion protein is expressed by calretinin neurons. Recordings were made at room temperature from YFP-negative neurons in lamina II to assess optically-evoked EPSCs (oEPSCs). These oEPSCs could result from 3 sources: (1) monosynaptic input from excitatory calretinin cells (eCRs), (2) polysynaptic circuits that include an eCR, or (3) a di-synaptic circuit in which an iCR forms an axoaxonic synapse on a primary afferent central terminal that is itself presynaptic to the recorded neuron (Fig 11a).

**Fig 11.**
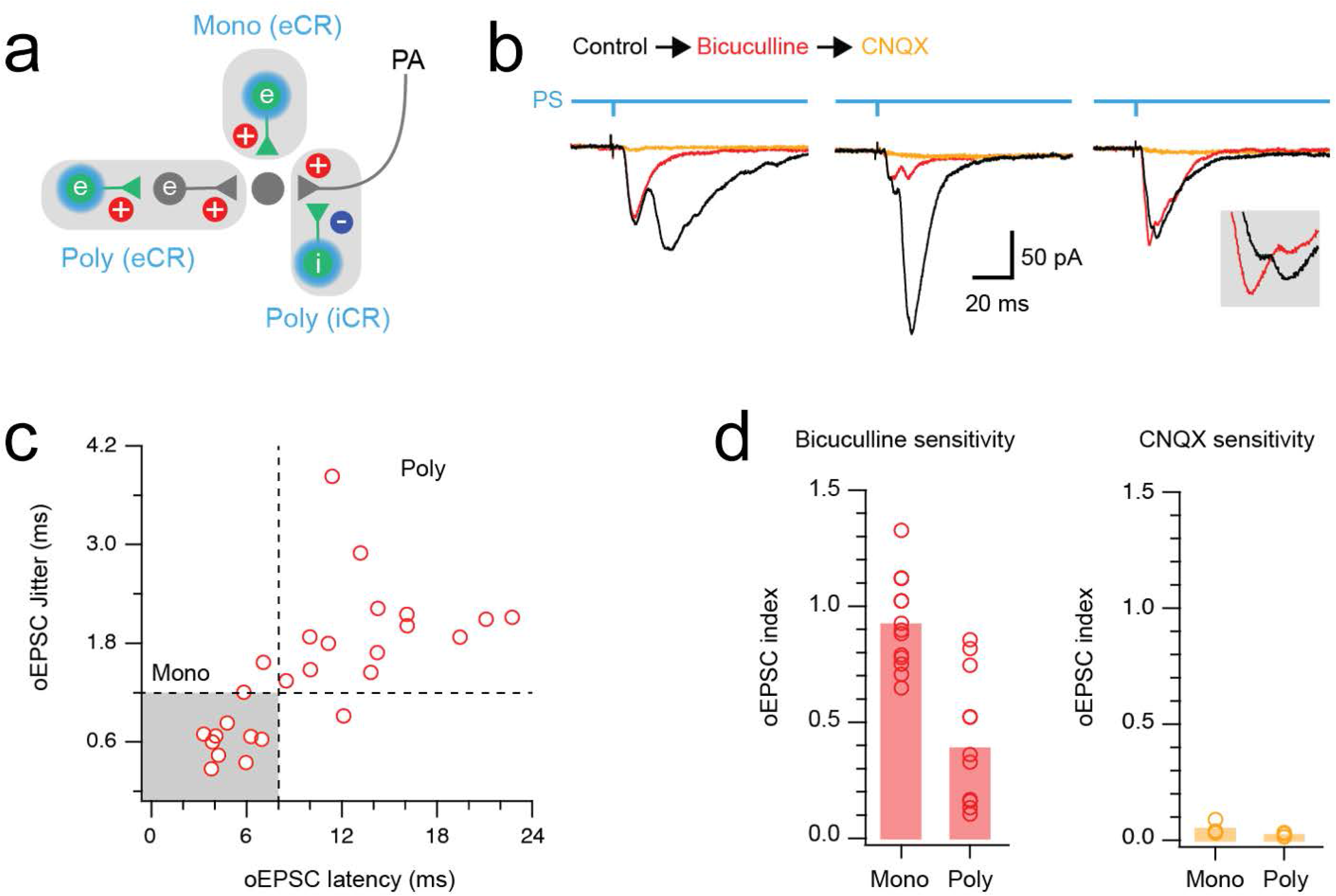
Functional identification of iCR-mediated presynaptic inhibition. **a:** schematic diagram summarising potential sources of excitatory input in response to photostimulation of channelrhodopsin-expressing calretinin neurons. Excitatory calretinin (eCR) neurons with direct input to a recorded cell give rise to a short latency monosynaptic input (Mono (eCR)). eCR neurons can also produce longer latency polysynaptic input (Poly (eCR)) by activating interposed excitatory interneurons. Alternatively, iCR neurons can elicit excitatory signals by releasing GABA onto primary afferents (PA), producing primary afferent depolarisation that can be observed as a longer latency polysynaptic excitatory input (Poly (iCR)). **b:** representative recordings show pharmacological dissection of optogenetic EPSCs (oEPSCs) during photostimulation in CR^Cre^;Ai32 mice (control = black, bicuculline = red, CNQX = orange). Left traces show a multicomponent oEPSC response (black trace) where bicuculline application abolished a longer latency component (red), while the short latency component was bicuculline resistant and CNQX sensitive. Middle traces show a multicomponent oEPSC response (black trace) where bicuculline dramatically reduced longer latency components, with the remaining oEPSCs abolished by CNQX. Right traces show a multicomponent oEPSC response (black trace) that exhibited little change on bicuculline application but was abolished when CNQX was applied. Inset shows expanded response peaks, highlighting a modest increase in the first, and decrease in the second peak after bicuculline addition. **c:** scatter plot shows oEPSC latency versus jitter for CR photostimulation-evoked oEPSC responses (14 recordings). A population of short latency (<8 ms) low jitter (<1.2 ms) oEPSCs are likely to result from direct monosynaptic input from eCRs (grey shading). In contrast, polysynaptic circuits driven by eCRs or iCRs produce oEPSCs with longer latencies and higher jitter. **d:** group data plots compare the sensitivity of monosynaptic and polysynaptic oEPSCs to bicuculline (left) and CNQX (right), using the oEPSC index. This is defined as the oEPSC amplitude in the presence of the drug divided by the oEPSC amplitude prior to drug application, with an oEPSC index of 1 indicating that the drug has no effect, and an index of 0 indicating that the drug completely blocks the oEPSC response. Several polysynaptic oEPSCs are reduced by bicuculline, whereas monosynaptic oEPSCs are resistant. All oEPSCs are CNQX sensitive.

Patch clamp recordings of multicomponent oEPSC responses, assessed under control conditions and following addition of bicuculline, were used to test for the contribution of iCR-evoked PAD inputs (n = 14) (Fig 11b). As expected, short latency oEPSC in these recordings were consistently bicuculline insensitive, and presumably resulted from monosynaptic input from eCRs. However, the response to bicuculline was more variable for longer latency oEPSCs ranging from complete block to minimal change. In a subset of bicuculline-resistant oEPSCs, subsequent CNQX addition abolished the remaining response (n = 5). Based on our past analyses^23, 25^, we define oEPSC currents with a latency of <8ms and jitter <1.2ms as monosynaptic (n = 11) and those with longer latencies and larger jitter values (n = 14) as polysynaptic (Figure 11c). Group comparisons of bicuculline sensitivity showed that monosynaptic responses were bicuculline-resistant (oEPSC index = 0.92 ± 0.19), whereas bicuculline significantly reduced polysynaptic oEPSC amplitude (oEPSC index = 0.39 ± 0.28, p = 0.001) (Fig 11d). All remaining responses were CNQX sensitive (oEPSC index = 0.05 ± 0.03 monosynaptic *vs.* 0.02 ± 0.01 polysynaptic, p = 0.145). Together, these results demonstrate that polysynaptic oEPSCs arise from a combination of eCR and iCR activation. The significant fall in polysynaptic oEPSC index in the presence of bicuculline provides functional evidence for axoaxonic synapses between iCRs and primary afferents. In contrast, a population of eCRs produce monosynaptic oEPSCs, as described in our previous work^69^.

## DISCUSSION

Our main findings are that: (1) the Rorb^CreERT2^ mouse line captures a subset of iCRs that appear to have reciprocal synaptic connections to the three main types of non-peptidergic nociceptors; (2) iCRs receive axodendritic synapses from central terminals of synaptic glomeruli in lamina II; (3) their axons form axoaxonic synapses onto these terminals, often as part of synaptic triads; and (4) optogenetic activation of iCRs leads to GABA-mediated oEPSCs in lamina II neurons. Taken together these findings suggest that the iCRs are a major postsynaptic target for non-peptidergic nociceptors and exert presynaptic inhibitory control over these afferents.

### Axoaxonic synapses and presynaptic inhibition

There is a wealth of evidence that GABAergic axoaxonic synapses on primary afferent terminals within the spinal cord generate presynaptic inhibition^7^. This is believed to operate through PAD^70^, which underlies the negative dorsal root potential, observed when an adjacent dorsal root is stimulated^71^. Two mechanisms have been proposed to link PAD to presynaptic inhibition: (1) blockade of action potential invasion into primary afferent terminals, or (2) a decrease in the amplitude of these action potentials, resulting in less Ca^2+^ entry into the terminals and reduced glutamate release^7^. An advantage of this form of inhibition over the postsynaptic inhibition generated by axodendritic and axosomatic synapses is that it is highly selective, being restricted to particular types of sensory input^7^.

In addition to non-peptidergic nociceptors, central terminals of several other types of primary afferent neuron receive axoaxonic synapses in the spinal cord. These include proprioceptive afferents, as well as A-LTMRs and C-LTMRs^72–74^. We have previously identified two other populations of inhibitory interneurons in the spinal cord that give rise to axoaxonic synapses on different types of primary afferent. We showed that central terminals of group 1a proprioceptive afferents in the ventral horn receive axoaxonic synapses exclusively from neurons located in the deep medial part of the dorsal horn that express the 65 kD isoform of glutamic acid decarboxylase (GAD65)^75^. These were later named GABApre neurons, and shown to generate presynaptic inhibition of Ia afferent terminals^68^. We subsequently identified PV-expressing cells as a source of axoaxonic synapses onto several different types of A-LTMR, including Aβ and Aδ hair follicle afferents, and those that innervate glabrous skin^22, 23, 46^. Until now, the origin of presynaptic inhibitory input to non-peptidergic nociceptors was not known. Here, we demonstrate that iCRs are a major source of axoaxonic synapses on the central terminals of each of the main classes of these afferents.

Although it is widely believed that GABAergic axoaxonic synapses mediate presynaptic inhibition, Hari et al^76^ recently suggested that GABA had a facilitatory effect on proprioceptive afferents, by preventing failure of action potentials to propagate through axonal branch points. This view was based partly on the apparent lack of GABA_A_ receptors on boutons formed by these afferents. However, Fink et al^68^ had previously demonstrated that optogenetic activation of GABApre neurons reduced the size of dorsal root-evoked EPSCs in motoneurons, thus providing direct evidence for presynaptic inhibition. Lorenzo et al^77^ have demonstrated expression of GABA_A_ receptor subunits in the central terminals of non-peptidergic afferents (identified by IB4 binding) in the rat, and here we show that activation of iCRs depolarises primary afferent terminals. In addition, we provide evidence that boutons of iCRs form both axoaxonic and axodendritic synapses, with at least some of the latter being in synaptic triads. It is therefore likely that the predominant role of iCRs is to generate presynaptic inhibition of the non-peptidergic nociceptor terminals, together with postsynaptic inhibition of their synaptic targets.

### Transgenic mouse lines for targeting iCRs

We initially used the Pnoc::GFP;CR^Cre^;Ai9 cross for electrophysiological studies, as this cross revealed over 75% of iCRs, and it was relatively straightforward to target cells that co-expressed tdTomato and GFP for electrophysiological recording. However, this cross was not suitable for anatomical studies of synaptic circuits, due to the high density of profiles that expressed each fluorescent protein. We therefore searched for a more selective approach for capturing these cells.

We noted that a Rorb^CreERT2^ mouse generated in a previous study^46^ could be used to label a small population of neurons located within the region innervated by non-peptidergic nociceptors, as revealed with IB4 binding. We found that these cells were largely restricted to the iCR population, although crosses involving the Rorb^CreERT2^ mouse only captured around 30% of these cells.

Crossing the Rorb^CreERT2^ mouse with various reporter lines allowed reliable targeting of iCRs for whole-cell recording and was also suitable for studies of synaptic circuitry, as the density of labelled profiles was far lower. Although our focus was on the Rorb cells in lamina II, it is of interest that Rorb is also expressed by the GABApre cells in the deep dorsal horn that generate presynaptic inhibition of proprioceptive afferents^78^.

In order to maximise the yield of iCRs for our EM studies, we turned to an intersectional strategy involving intraspinal injection of an AAV that depended on co-expression of Cre and Flp to deliver GFP to CR^Cre^;VGAT^Flp^ mice. This approach should be efficient and reliable, and this was borne out by the distribution of GFP-expressing cells. These were largely restricted to the SDH, and had processes that remained within this region, consistent with the islet morphology of iCRs. Although this approach requires surgery to deliver the virus, it has the advantage of being highly flexible. For example, we were able to restrict expression to the Gaba9/DI1 population by using Tac1^Cre^;VGAT^Flp^ mice, and it might be possible to target the Gaba8 population of Haring et al by using mice in which Cre is under transcriptional control of the gene for keratin 17 (Krt17), which is highly expressed in this population^18^. This would then allow a comparison of the synaptic circuits engaged by these two subsets of iCRs. In addition, this approach will allow investigation of the functions of these cells through chemogenetic or optogenetic manipulation.

### Functional roles of iCRs

Our anatomical data suggest that the iCRs receive a powerful excitatory synaptic input from non-peptidergic nociceptors. This is consistent with the finding that most of the cells for which we could record primary afferent input showed monosynaptic C fibre responses, and that this was often sufficient to drive action potential firing. TRPV1 is expressed by afferents belonging to the NP2 and NP3 classes, but not those of NP1,^9, 51, 52, 56, 79, 80^, and although most of the cells tested showed an increase in mEPSC frequency in the presence of capsaicin, application of capsaicin had only minor effects on dorsal root-evoked EPSCs. This suggests that these cells receive a major input from TRPV1-negative afferents, which are presumably those belonging to NP1, with a smaller contribution from those of the NP2 and NP3 populations. However, it is also possible that some of the input from TRPV1-expressing afferents originates from peptidergic nociceptors. We also provide evidence that the iCRs preferentially target the same types of afferent, and we show that they frequently give rise to axoaxonic synapses in lamina II. The high proportion (70%) of Rorb-derived boutons that target MrgD (NP1) afferents may reflect the fact that these are much more numerous than those belonging to NP2 and NP3. Alternatively, it might result from a preferential association between NP1 afferents and the subset of iCRs that are captured in the Rorb mouse. Most Rorb-derived boutons are associated with gephyrin puncta, suggesting that they formed axodendritic synapses, and we were able to identify synaptic triads, in which the iCR axon was presynaptic to a glomerular central terminal and also to a dendrite that was itself innervated by the central terminal. This would allow for both presynaptic and postsynaptic inhibition of the afferent input to other dorsal horn neurons (Fig 12). The iCRs therefore appear to have a highly selective pattern of both synaptic input and output, in that they are closely associated with non-peptidergic nociceptors and their postsynaptic targets. This selectivity is reflected in their islet morphology, since both their axonal and dendritic arbors are largely restricted to the band within lamina II that receives input from these afferents. A similar arrangement applies to the PV-expressing inhibitory interneurons in laminae IIi-III, many of which are also islet cells, except that in this case they are associated with A-LTMRs, and are therefore located more ventrally. Many other dorsal horn neurons have dendrites and axons that are far more extensive in the dorsoventral axis^81–83^, and this is likely to reflect a more diverse pattern of synaptic inputs to and outputs from these cells.

**Fig 12.**
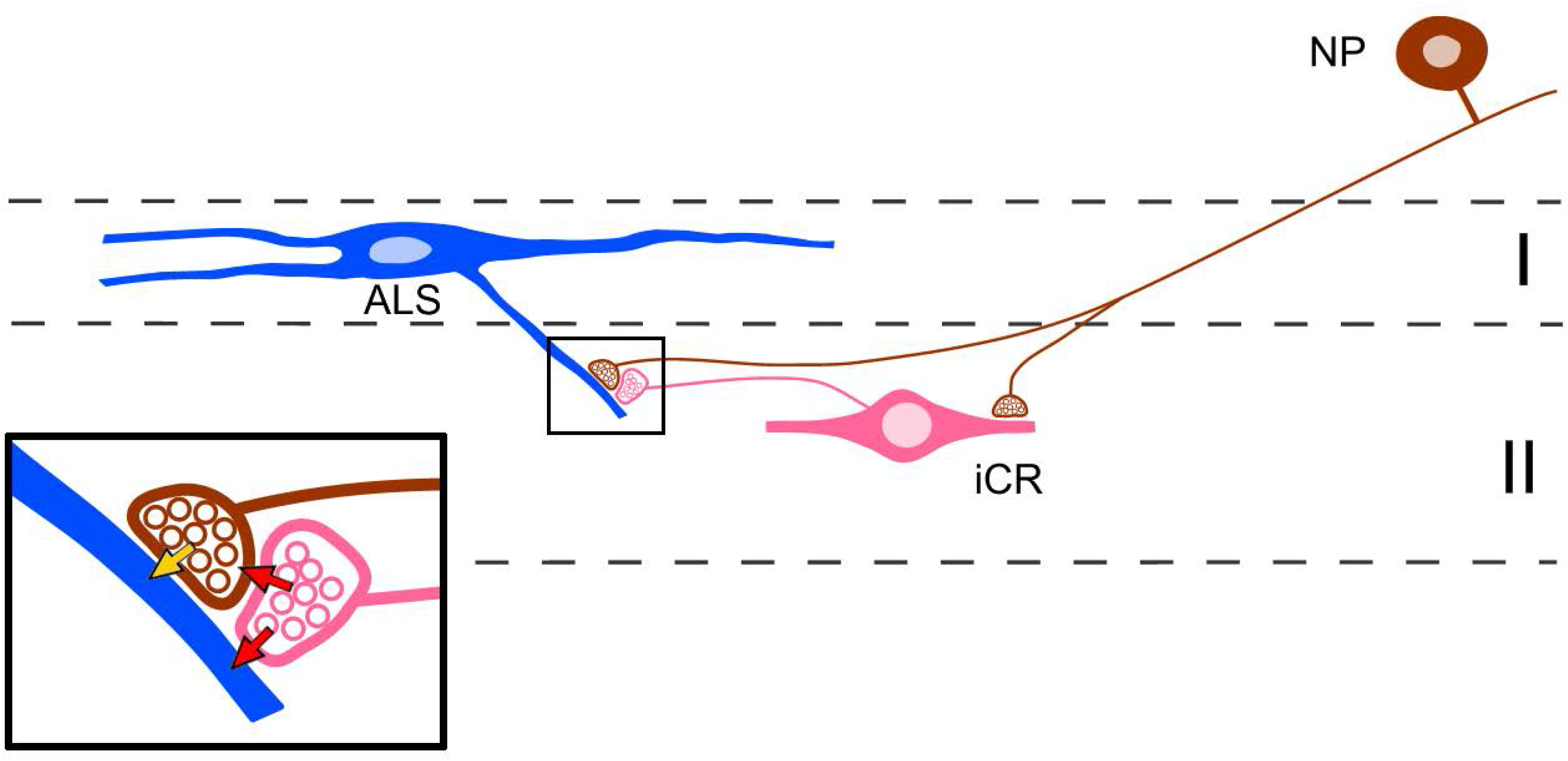
Schematic diagram summarising synaptic connections involving inhibitory calretinin cells and primary afferents. A non-peptidergic nociceptor (NP) forms excitatory synapses onto an inhibitory calretinin cell (iCR) in lamina II and a lamina I projection neuron belonging to the anterolateral system (ALS). The ALS cell is shown as representative of spinal neurons that are innervated by NP afferents. The iCR axon contributes to a synaptic triad, which is shown in more detail in the inset. The excitatory (glutamatergic) synapse between the NP afferent and the ALS cell is indicated with a yellow arrow. The axon of the iCR is presynaptic to the NP afferent (upper red arrow) at an axoaxonic synapse, and to the ALS cell dendrite (lower red arrow) at an axodendritic synapse. These synapses are both GABAergic, and mediate pre-and postsynaptic inhibition, respectively. GABA acting at the axoaxonic synapse will reduce glutamate release at the synapse from the NP afferent to the ALS cell (feedback inhibition), while at the axodendritic synapse it will directly inhibit the ALS cell (feedforward inhibition).

The roles of the iCRs in somatosensory processing and behaviour will therefore be determined both by the response properties of the non-peptidergic nociceptors with which they are associated, and also by the other types of dorsal horn neuron that receive input from these nociceptors. Each of the main NP classes contains cells that respond to both mechanical and thermal (heat) stimuli^52, 56, 84^. These responses are relatively modest for innocuous stimuli, but are enhanced for stimuli in the noxious range^56^, suggesting that NP afferents are able to encode stimuli that would be perceived as painful. However, despite this, recent studies have demonstrated that optogenetic activation of MrgD (NP1) afferents *in vivo* does not result in pain-related behaviours^85–87^. In addition the NP2 and NP3 classes express many itch-associated receptors^9, 56^ and both have been shown to function as pruritoceptors^26, 52^. Relatively little is yet known about other dorsal horn neurons that are innervated by these afferents; however, these include lamina I projection neurons belonging to the anterolateral system (ALS)^88^, and a population of excitatory interneurons that are defined by expression of GFP in a BAC transgenic GRP::GFP line^58^, and are thought to function as “secondary pruritoceptors”^79^.

The axoaxonic and axodendritic synapses formed by iCRs give rise to feedback (axoaxonic) and feedforward (axodendritic) GABAergic inhibition of the input from non-peptidergic nociceptors to other dorsal horn neurons, presumably including lamina I ALS projection neurons^88^ (Fig 12). This inhibition would be triggered whenever these afferents are activated, and this may partly explain why optogenetic activation of MrgD afferents in otherwise healthy mice is not perceived as painful^85–87^. Importantly, activation of these afferents was apparently perceived as painful in models of both neuropathic^87^ and inflammatory^85^ pain, suggesting that they contribute to persistent pain states. The iCRs may therefore be an attractive therapeutic target for the treatment of pathological pain and itch.

## METHODS

### Animals

All experiments were approved by the Ethical Review Process Applications Panel of the University of Glasgow or by the Animal Care and Ethics Committee at the University of Newcastle. Experiments undertaken in Glasgow were performed in accordance with the European Community directive 86/609/EC and the UK Animals (Scientific Procedures) Act 1986. The study was carried out in compliance with the ARRIVE guidelines. A list of the mouse lines used is provided in Table S1. Crosses between these lines were generated as described in Results. Mice of both sexes aged between 4 and 29 weeks were used in the study. For crosses involving the Rorb^CreERT2^ line, animals received intraperitoneal injections of tamoxifen (1 mg) between the ages of P14-P18.

### Anatomical analysis of Neurobiotin-filled calretinin cells

Five eGFP-expressing cells that displayed tonic-firing patterns were identified in whole-cell patch-clamp experiments involving parasagittal slices from 3 CR::GFP mice (2 female, 1 male), performed as described previously^34, 36^. At the end of the recording session, the slices were fixed by immersion in 2.5% glutaraldehyde in 0.1M PB for at least 24 hours. They were then re-sectioned at 60 µm thickness on a Leica VT1000 vibrating blade microtome. Sections were treated with 0.03% H_2_O_2_ for 30 minutes to block endogenous peroxidase activity and incubated in Avidin-peroxidase (1:1000 in PB; Sigma-Aldrich, UK) for 24 hours to reveal the Neurobiotin present in the recorded cells. Peroxidase labelling was visualised with 3,3’,5,5’-diaminobenzidine (DAB; Sigma-Aldrich), and the sections were osmicated, block stained with uranyl acetate and flat-embedded in Durcupan resin as described previously^22^. Complete reconstructions of the cell body, together with dendritic and axonal arbors were made for each cell, using a Nikon Optiphot-2 microscope with a 100× oil immersion lens (NA 1.4) and drawing tube. Individual sections containing parts of the axon and dendritic tree from each cell were mounted onto resin stubs. Serial ultrathin sections were then cut from these blocks with a Reichert Ultracut S (Leica), collected onto Formvar-coated copper slot grids and stained with lead citrate. These were viewed using a Philips CM100 transmission electron microscope.

### Immunofluorescence staining, confocal microscopy and analysis

Multiple-labelling immunohistochemistry was performed as described previously^25, 83^ on 60 μm thick transverse or sagittal sections from the L2-L5 segments, which were cut with vibrating blade microtomes (Leica VT1200 or VT1000). Sources and concentrations of antibodies are listed in Table S2. Sections were incubated for 3-5 days at 4°C in mixtures of primary antibodies diluted in PBS that contained 0.3 M NaCl, 0.3% Triton X-100 and 5% normal donkey serum, and then overnight in species-specific secondary antibodies (Jackson Immunoresearch, West Grove, PA, USA), which were raised in donkey and conjugated to Alexa488, Alexa647, Rhodamine Red, Pacific Blue or biotin. All secondary antibodies were diluted 1:500 in the same diluent, apart from those conjugated to Rhodamine Red, which were diluted 1:100. Biotinylated antibodies were revealed with Pacific Blue conjugated to avidin (1:1,000; Life Technologies, Paisley, UK). In some cases sections were incubated in biotinylated IB4 (1:500; Sigma, RRID:AB_2313663), which was revealed with avidin-Pacific Blue. Following immunoreaction, sections were mounted in anti-fade medium and stored at -20°C. They were scanned with either a Zeiss LSM710 confocal microscope with Argon multi-line, 405 nm diode, 561 nm solid state and 633 nm HeNe lasers, or with a Zeiss LSM900 Airyscan confocal microscope with 405, 488, 561 and 640 nm diode lasers. Confocal image stacks were obtained through a 20× dry lens (numerical aperture, NA, 0.8), or through 40× (NA 1.3) or 63× (NA 1.4) oil immersion lenses with the confocal aperture set to 1 Airy unit or less. All analyses were performed with Neurolucida for Confocal software (MBF Bioscience, Williston, VT, USA).

To determine the efficiency of capture of iCRs in different genetic crosses, we analysed tissue from 3 Pnoc::GFP;CR^Cre^;Ai9 mice (all male) and from 5 Rorb^CreERT2^;Ai9 mice (3 male, 2 female). Quantification of synaptic inputs and outputs of the Rorb cells was performed on tissue from 3 Rorb^CreERT2^;Ai9;MrgD^ChR2-YFP^ mice (2 male, 1 female), 3 Rorb^CreERT2^;Ai9 mice (2 male, 1 female), 2 Rorb^CreERT2^;Ai34;MrgD^ChR2-YFP^ mice (1 male, 1 female), and 2 Rorb^CreERT2^;Ai34 mice (1 male, 1 female).

### Fluorescent *in situ* hybridisation

Multiple-labelling FISH was performed with RNAscope probes and RNAscope fluorescent multiplex reagent kit 320850 (ACD BioTechne; Newark, CA 94560). Fresh frozen lumbar spinal cords were rapidly removed from 3 Rorb^CreERT2^;Ai9 mice (2 male, 1 female) and 3 wild-type mice (2 male, 1 female), embedded in OCT mounting medium and cut into 12 μm thick transverse sections with a cryostat (Leica CM1950; Leica; Milton Keynes, UK). These were mounted in Prolong-Glass anti-fade medium with NucBlue (P36981; Invitrogen, Thermofisher Scientific) non-sequentially onto Superfrost Plus slides (48311703, VWR, Lutterworth, UK) so that sections on the same slide were at least 40 μm apart in the z plane. Reactions were carried out according to the manufacturer’s recommended protocol. The probes used in this study, and the proteins that they correspond to are listed in Table S3. Sections from the Rorb^CreERT2^;Ai9 mice were incubated in probes directed against *tdTomato*, *Gad1* and *Tac1*, while those from wild-type mice were incubated in probes against *Slc32a1*, *Tac1* and *Pam*. Positive and negative control probes were also tested on other sections (Table S3). Sections were scanned through the full thickness with a 1 μm z-step on the LSM710 confocal microscope.

### Electrophysiological characterisation of iCRs

Electrophysiological recordings were performed on spinal cord slices from a total of 78 mice, which included 33 Pnoc::GFP;CR^Cre^;Ai9 (20 female, 13 male), 39 Rorb^CreERT2^;Ai9 (24 female, 15 male) and 6 Rorb^CreERT2^;Ai32 mice (3 female, 3 male). Recordings were made from a total of 200 cells, that included 82 (47 female, 35 male) from Pnoc::GFP;CR^Cre^;Ai9 mice, 107 (61 female, 46 male) from Rorb^CreERT2^;Ai9 mice and 11 (6 female, 5 male) from Rorb^CreERT2^;Ai32 mice.

Spinal cord slices were prepared as described previously^82, 83^. Mice were anaesthetised with isoflurane, decapitated and the spinal cord (in some cases with dorsal roots attached) was removed by performing a ventral laminectomy in ice-cold dissection solution. In some cases the mouse was terminally anaesthetised with pentobarbital (20mg i.p.) followed by transcardial perfusion with ice-cold dissection solution, prior to decapitation. The lumbar spinal cord, in some cases with either L3 and L4, or L4 and L5 dorsal roots attached, was embedded in low-melting point agar (∼3%; Thermo Fisher Scientific) and parasagittal slices were cut with a vibrating blade microtome (Thermo Scientific, Microm HM 650V), at a thickness of 300 µm (no roots) or 400-500 µm (roots attached). Slices were allowed to recover in recording solution for at least 30 minutes at room temperature. In some cases slices were placed in a *N*-methyl-D-glucamine (NMDG)-based recovery solution for 15 minutes at ∼32°C before being placed in a modified recording solution at room temperature for at least 30 minutes. The solutions used contained the following (in mM); dissection, 251.6 sucrose, 3.0 KCl, 1.2 NaH_2_PO_4_, 0.5 CaCl_2_, 7.0 MgCl_2_, 26.0 NaHCO_3_, 15.0 glucose; NMDG recovery, 93.0 NMDG, 2.5 KCl, 1.2 NaH_2_PO_4_, 0.5 CaCl_2_, 10.0 MgSO_4_, 30.0 NaHCO_3_, 25.0 glucose, 5.0 Na-ascorbate, 2.0 thiourea, 3.0 Na-pyruvate, and 20.0 HEPES; modified recording, 92.0 NaCl, 2.5 KCl, 1.2 NaH_2_PO_4_, 2.0 CaCl_2_, 2.0 MgSO_4_, 30.0 NaHCO_3_, 25.0 glucose, 5.0 Na-ascorbate, 2.0 thiourea, 3.0 Na-pyruvate, and 20.0 HEPES; and recording, 125.8 NaCl, 3.0 KCl, 1.2 NaH_2_PO_4_, 2.4 CaCl_2_, 1.3 MgCl_2_, 26.0 NaHCO_3_, 15.0 glucose. All solutions were bubbled with 95% O_2_/5% CO_2_.

Neurons were visualised using a fixed stage upright microscope (BX51; Olympus, Southend-on-Sea, UK) equipped with a 40× water immersion objective, infrared differential interference contrast (IR-DiC) illumination, and a CCD camera (QImaging Retiga Electro; Teledyne Photometrics, Birmingham, United Kingdom). Fluorescently labelled cells were visualised using a 470 nm or 550 nm LED (pE-100; CoolLED, Andover, United Kingdom).

Targeted whole-cell patch-clamp recordings were made from cells in lamina II of the dorsal horn that expressed both GFP and tdTom (Pnoc::GFP;CR^Cre^;Ai9), or expressed only tdTom (Rorb^CreERT2^;Ai9) or YFP (Rorb^CreERT2^;Ai32). Recordings were made at room temperature, using patch pipettes that had resistances of 3-7 MΩ when filled with internal solution that usually contained (in mM); 130 K-gluconate, 10 KCl, 2.0 MgCl_2_, 10 HEPES, 0.5 EGTA, 2 ATP-Na_2_, 0.5 GTP-Na, and 0.2% Neurobiotin, pH adjusted to 7.3 with 1.0M KOH. In some experiments that involved dorsal root stimulation, a Cs-based intracellular solution containing the following (in mM) was used; 120 Cs-methylsulfonate, 10 Na-methylsulfonate, 10 EGTA, 1 CaCl_2_, 10 HEPES, 5 N-(2,6-dimethylphenylcarbamoylmethyl) triethylammonium chloride (QX-314-Cl), 2 Mg_2_-ATP, and 0.2% Neurobiotin, pH adjusted to 7.2 with 1.0M CsOH. Data were recorded and acquired with a Multiclamp 700B amplifier and pClamp 10 software (both Molecular Devices), and were filtered at 4 kHz and digitised at 10 kHz.

After stable whole-cell configuration was achieved, the cells were voltage clamped at -60 mV and a series of 100 ms voltage steps from -70 to -50 mV, in 2.5 mV increments, were delivered to determine the I-V relationship, which was used to calculate the resting membrane potential (RMP) and input resistance. Any cell that had a RMP less negative than -40 mV was excluded from all analysis.

Action potential firing patterns were assessed by applying 1 s depolarising current steps, that increased in amplitude by 5 pA per sweep, from a membrane potential of around -60 mV. Firing patterns were classified on the basis of previously published criteria^16, 34, 38, 81, 89–91^. Cells were classified as tonic firing if a continuous action potential discharge occurred throughout the depolarising step; transient firing if the action potential discharge occurred only during the early phase of the current step; and single spike if only 1 or 2 action potentials occurred at the start of the step.

The incidence of subthreshold voltage-activated currents was investigated by applying a voltage step protocol in which cells were stepped from -60 mV to -90 mV for 1 s and then to -40 mV for 200 ms^34, 92^, with automated leak subtraction used to remove capacitive and leak currents. This protocol enables identification of two types of inward current and two types of transient outward current. During the hyperpolarising step from -60 to -90 mV, a slow inward current, that is consistent with the hyperpolarisation-activated (I_h_) current, can be observed. During the depolarising step from -90 to -40 mV three types of current can be observed. A transient inward current is considered to reflect low-threshold ‘T-type’ calcium current (I_Ca,T_). The two transient outward currents are consistent with A-type potassium currents (I_A_) and are defined as rapid (I_Ar_) or slow (I_As_) on the basis of their kinetics. The amplitude of I_h_ currents was measured during the final 200 ms of the hyperpolarising step, and I_Ca,T_ amplitude was measured as the peak of the transient inward current.

Primary afferent input to iCRs was assessed by recording evoked excitatory postsynaptic currents (eEPSCs) in spinal cord slices with dorsal roots attached. Cells were voltage clamped at -70 mV and the dorsal root stimulated using a suction electrode connected to an ISO-Flex stimulus isolator (AMPI, Jerusalem, Israel). In most cases the slice had two dorsal roots attached^81, 82, 93^, either L3 and L4, or L4 and L5, and each was stimulated independently using a separate suction electrode and stimulus isolator^80, 81, 92^. To characterise the primary afferent input to cells, the dorsal root was stimulated three times at low-frequency (0.05 Hz, 0.1 ms duration), using the following intensities to activate Aβ, Aδ and C fibres; 25 µA for Aβ, 100 µA for Aδ and 1mA for C fibres. Primary afferent input was characterised as monosynaptic or polysynaptic by stimulating the dorsal roots twenty times using the following intensities and frequencies; Aβ, 25 µA/20 Hz; Aδ, 100 µA/2 Hz; and C, 1mA/1 Hz. A fibre responses were considered monosynaptic if there was an absence of response failures and the response latency varied by ≤2 ms, whereas C fibre responses were classified as monosynaptic if there was an absence of failures, regardless of whether the response latency was variable^94^. The estimated conduction velocity of monosynaptic responses was calculated using the response latency, measured as the time between the stimulus artefact and the onset of the eEPSC, and the length of the stimulated dorsal root, measured as the distance between the suction electrode and the dorsal root entry zone. To determine whether monosynaptic C fibre input to iCRs was sufficient to drive action potential firing, dorsal roots were stimulated three times at 0.05 Hz (0.1 ms duration) at an intensity of 1 mA during current clamp recordings, at a membrane potential of around -60 mV. This was performed for a subset of the cells that were found to have monosynaptic C fibre input.

The effect of activating TRPV1 channels on the monosynaptic C fibre input was assessed in a subset of cells. Monosynaptic C fibre eEPSCs were evoked at 1 mA (0.05 Hz, 0.1 ms duration) for 10 minutes (baseline) followed by a further 10 minutes in the presence of the TRPV1 agonist capsaicin (2 µM), which was added to the recording solution and washed into the bath. Inputs were classified as ‘sensitive’ if capsaicin reduced the ratio of the mean eEPSC peak amplitude recorded in the final 5 minutes of application to 75% of the mean peak amplitude recorded in the final 5 minutes of baseline, and ‘insensitive’ if this threshold was not met. The incidence of input from TRPV1-expressing primary afferents was also assessed in a separate group of cells by recording miniature EPSCs (mEPSCs), in the presence of TTX (0.5 µM), bicuculline (10 µM) and strychnine (1 µM), at a holding potential of -70 mV. Baseline mEPSCs were recorded for 5 minutes, followed by a further 5 minutes in the presence of capsaicin (2 µM). The final 2 minutes of baseline and capsaicin recordings were analysed using MiniAnalysis (Synaptosoft). Events were automatically detected by the software and were then accepted or rejected after visual examination. Cells were classified as receiving input from TRPV1-expressing afferents if capsaicin application caused a significant leftward shift in the distribution of mEPSC inter-event intervals.

The response of CR islet cells to opiates was investigated by voltage clamping cells at -60 mV, in the presence of TTX (0.5 µM), bicuculline (10 µM) and strychnine (1 µM), and bath applying one of the following: the µ-opioid receptor agonist DAMGO (3 µM), the δ-opioid receptor agonist [D-Ala^2^] Deltorphin II (1 µM) or the κ-opioid receptor agonist U69593 (1 µM). Following a baseline recording period of 4 minutes the agonist was washed into the bath for 5 minutes. To assess whether cells responded to these agonists mean current was measured in 30 s bins. Responses were compared to a 30 s baseline period, with cells being classified as responsive if a change in mean current of ≥5 pA was observed, and unresponsive if this threshold was not reached.

All chemicals were obtained from Sigma except: TTX, QX-314-Cl (Alomone, Jerusalem, Israel), bicuculline, DAMGO (Tocris, Abingdon, UK), Deltorphin II (Abcam, Cambridge, UK), sucrose, glucose, KCl (VWR, Lutterworth, UK), NaH_2_PO_4_ (VWR, Lutterworth, UK), sucrose, glucose, agarose (ThermoFischerScientific) and Neurobiotin (Vector Labs Peterborough, UK).

### Intersectional targeting with AAV.Con/Fon.GFP

Four female CR^Cre^;VGAT^Flp^ mice received injections of an AAV that expressed GFP in the presence of both Cre and Flp recombinases under transcriptional control of the human elongation factor 1 alpha promoter fragment hEF1α (AAV1.C^on^/F^on^.GFP) into the right side of the spinal cord as described previously^26, 82^. Briefly, the mice were anaesthetised with isoflurane and received injections in the spaces between the T12/T13 and T13/L1 vertebrae, to allow targeting of the L3 and L5 spinal segments. In some cases an additional injection was made through a small hole drilled in the lamina of T13 to allow targeting of the L4 segment. At each site, 300 nl of virus containing 8.1 × 10^8^ GC was injected 400 μm lateral to the midline and 300 μm below the pial surface. The wound was closed and the mice received perioperative analgesia (0.5 mg/kg buprenorphine and 5 mg/kg carprofen). After a 13-23 day survival period the mice were reanaesthetised and perfused transcardially with a fixative that contained 4% formaldehyde. In two cases (used for electron microscopy) the fixative also contained 0.2% glutaraldehyde.

Tissue from the mice fixed with formaldehyde/glutaraldehyde was processed for immunoperoxidase labelling. The L4 segments were fixed overnight at 4°C and cut into 60 μm thick transverse sections with a vibrating blade microtome (Leica VT1200). The sections were incubated for 3 days in rabbit anti-GFP (see Table S2), followed by overnight incubations in biotinylated donkey anti-rabbit IgG and avidin-HRP. All of these reagents were diluted in PBS. The sections were then reacted with DAB, and processed for electron microscopy. We used the same procedure as described above, except that post-fixation was performed with a “reduced osmium” protocol as described by Zhang et al^66^. Ultrathin sections were cut onto Formvar-coated slot grids and stained with lead citrate, before being viewed on the EM. Tissue from the other 2 mice was cut into transverse or sagittal sections, which were reacted to reveal GFP with an immunofluorescence method, as described above.

Two male Tac1^Cre^;VGAT^Flp^ mice received intraspinal injections of AAV1.C^on^/F^on^.GFP into the L3 and L5 segments on the right side as described above. In this case 1.6 × 10^9^ GC in 300 nl was injected at each site. After a 7 day survival time, mice were perfused with fixative containing 4% formaldehyde. Spinal cord regions that included the injection sites was cut into transverse or sagittal sections, which were immunoreacted with antibody against GFP.

### Electrophysiological evidence for GABAergic transmission at axoaxonic synapses

Spinal cord slices were prepared from 11 CR^cre^;Ai32 mice (4 female, 7 male) for patch clamp recording, as previously described^69^. Briefly, mice were anaesthetised, the vertebral column isolated and submerged in carbogenated (95% O_2_ and 5% CO_2_) sucrose substituted artificial cerebrospinal fluid (sACSF) containing (in mM): 250 sucrose, 25 NaHCO3, 10 glucose, 2.5 KCl, 1 NaH2PO4, 1 MgCl2 and 2.5 CaCl2). The spinal cord was extracted using a ventral approach and sectioned (250μm sagittal slices) using a vibrating microtome (Leica VT-1000S, Heidelberg, Germany). All slices were transferred to an interface chamber containing carbogenated ACSF (118 mM NaCl substituted for sucrose) and allowed to stabilize for 1 hr at room temperature (22-24°C) before recordings commenced. Slices were transferred to a recording bath and superfused with carbogenated ACSF to achieve a pH of 7.3–7.4. Recordings were obtained from unidentified neurons in the superficial dorsal horn (laminae I-II) at room temperature (21–24°C), visualized with near-infrared differential interference contrast optics connected to an IR-sensitive camera (Jenoptik ProgRes MF cool).

Patch clamp recordings used a potassium gluconate-based internal solution containing (in mM): 135 C_6_H_11_KO_7_, 6 NaCl, 2 MgCl_2_, 10 HEPES, 0.1 EGTA, 2 MgATP, 0.3 NaGTP, pH 7.3 (with KOH) and were established in voltage clamp mode. Photostimulation was delivered using a CoolLED pE excitation system through the microscope light path at suprathreshold intensity (16 mW) with a duration of 1 ms (controlled by transistor-transistor logic pulses). Previous work in CR^cre^;Ai32 spinal slices has confirmed that this reliably recruits single action potential spikes in ChR2-expressing neurons^69^. Only recordings with multicomponent photostimulation-evoked excitatory synaptic responses (inward currents) were retained for inclusion in the study.

Pharmacology was then used to assess the nature of circuits responsible for these inputs. Bicuculline (10μM) was first applied to identify responses that relied on GABAergic signalling. We have previously demonstrated that bicuculline-sensitive responses arise when inhibitory ChR2-expressing axoaxonic input produces PAD, resulting in delayed excitatory responses in recorded postsynaptic neurons^23, 25^. If photostimulation responses remained, CNQX (10μM) was applied to abolish any remaining excitatory current.

Optically-evoked postsynaptic current (oEPSC) amplitude was measured from baseline just prior to photostimulation. The peak amplitude of responses was calculated from the average of 10 successive trials. The latency of oEPSCs was measured as the time between photostimulation and the evoked current onset. Jitter was defined as the standard deviation in latency of 10 successive trials. These properties were used to differentiate monosynaptic and polysynaptic responses, with the monosynaptic inputs exhibiting shorter latencies and minimal jitter. Changes in photostimulation-evoked postsynaptic current amplitude induced by drug application were used to calculate an oEPSC index. This index was defined as the oEPSC amplitude in the presence of the drug divided by the oEPSC amplitude prior to drug application.

### Statistical analysis

Electrophysiological differences between sexes or between recordings performed in Pnoc::GFP;CR^Cre^;Ai9 and Rorb^CreERT2^ tissue were compared using a t-test or Mann-Whitney U test. Capsaicin induced changes in the distribution of mEPSC inter-event intervals within individual cells were compared using a Kolmogorov-Smirnov 2-sample test and changes in mEPSC frequency in the whole population, between baseline and capsaicin recordings, were compared using a Wilcoxon test. Mann-Whitney U tests were used to compare transcript numbers in FISH experiments, while unpaired t-tests were used to compare bicuculline/CNQX sensitivity of monosynaptic and polysynaptic oEPSC amplitudes in optogenetic experiments.

## ACKNOWLEDGEMENTS

We thank Robert Kerr, Christine Watt and Iain Plenderleith for expert technical assistance, and Dr P. Ciofi for the gift of somatostatin antibody. Financial support from the Biotechnology and Biological Sciences Research Council (Grant numbers BB/J000620/1, BB/P007996/1 and BB/X000338/1), the National Health and Medical Research Council (NHMRC) of Australia (Grant numbers 631000 and 1043933), the Wellcome Trust (Grant numbers 102645/Z/13/Z and 219433/Z/19/Z), the Medical Research Council (Grant numbers MR/S002987/1 and MR/V033638/1), and the National Institute of Health (NIH; NS097344-08) is gratefully acknowledged. DDG is an investigator of the Howard Hughes Medical Institute.

## AUTHOR CONTRIBUTION

R.J.C., B.A.G., A.J.T. and D.I.H. conceived the project and designed experiments; O.C.D., A.C.D., M.B.M., K.A.B., T.J.B., M.A.G., K.M.S, E.P., A.M.B., B.A.G., A.J.T., and D.I.H. performed experiments; O.C.D., A.C.D., M.B.M., K.A.B., T.J.B., M.A.G., K.M.S., A.M.B., E.K., B.A.G., A.J.T. and D.I.H. analysed data; M.W., H.W., H.U.Z., D.D.G. provided reagents/animals; A.J.T. and D.I.H. wrote the paper; all of the authors provided feedback and contributed to the editing of the manuscript.

## DATA AVAILABILITY

The datasets generated and analysed during the current study are available from the corresponding authors on reasonable request.

