## Supplementary material for "Calretinin-expressing islet cells: a source of pre- and post-synaptic inhibition of non-peptidergic nociceptor input to the mouse spinal cord"

Number of words in abstract: 170
main text: 6591
online methods: 3423
Number of references: 94
Number of figures: 12 (plus 4 supplementary)

*Corresponding authors:

David I. Hughes, Spinal Cord Group, Sir James Black Building, University of Glasgow, Glasgow, G12 8QQ, UK.

Andrew J. Todd, Spinal Cord Group, Sir James Black Building, University of Glasgow, Glasgow, G12 8QQ, UK.

Brett A. Graham, School of Biomedical Sciences and Pharmacy, Faculty of Health and Medicine, University of Newcastle, Callaghan, NSW, Australia.

Fig S1

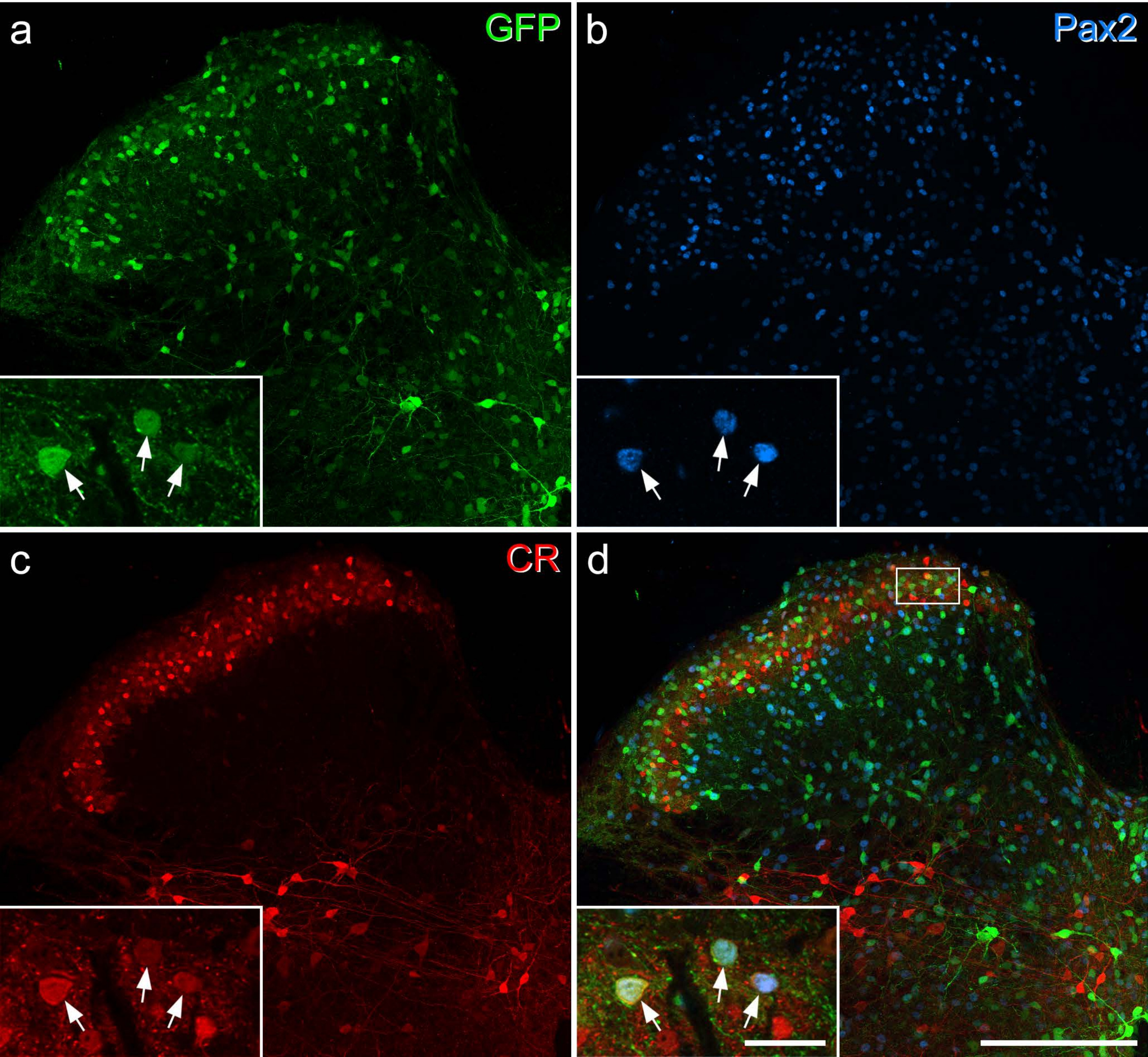

Fig S2

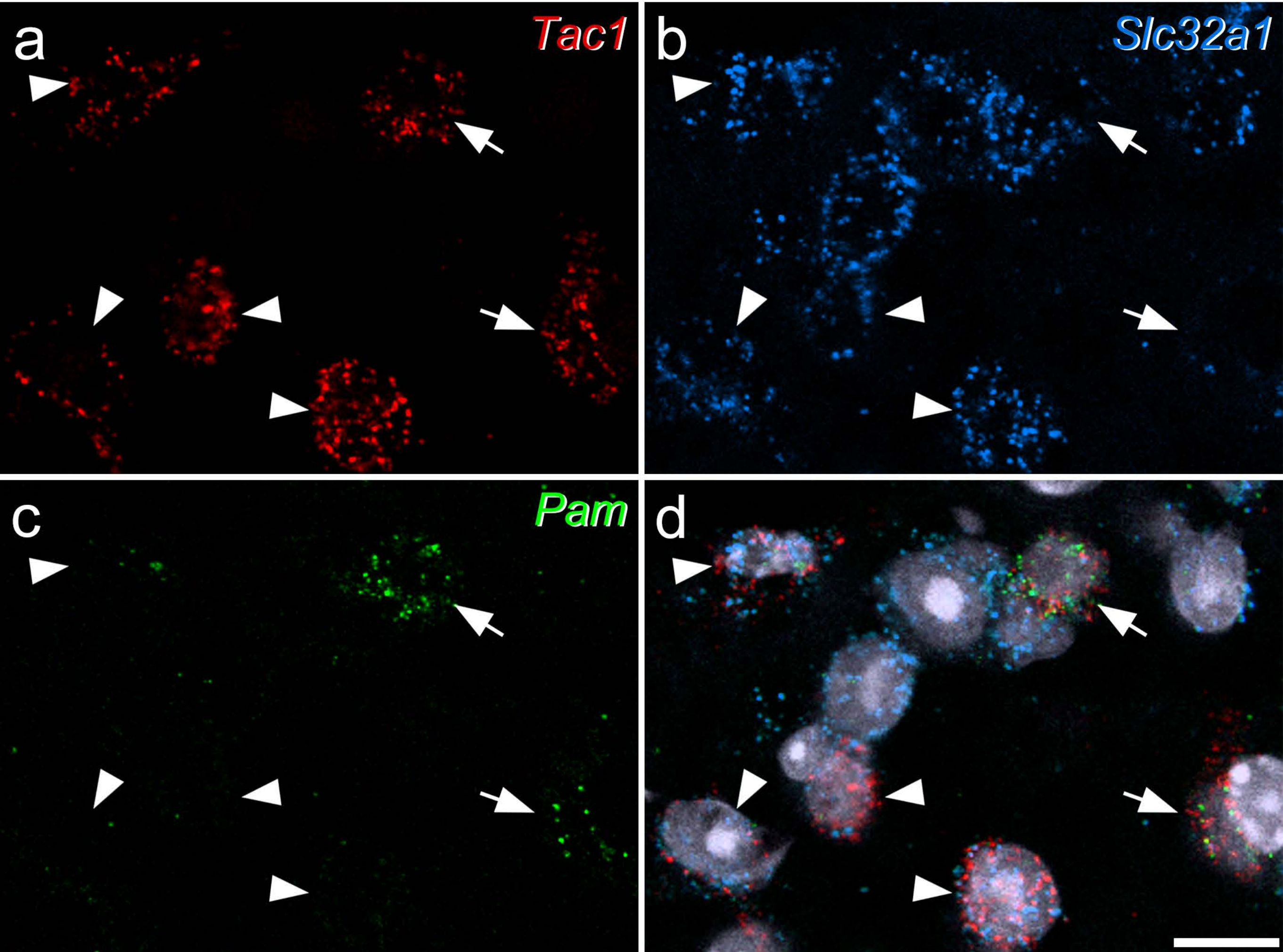

Fig S3

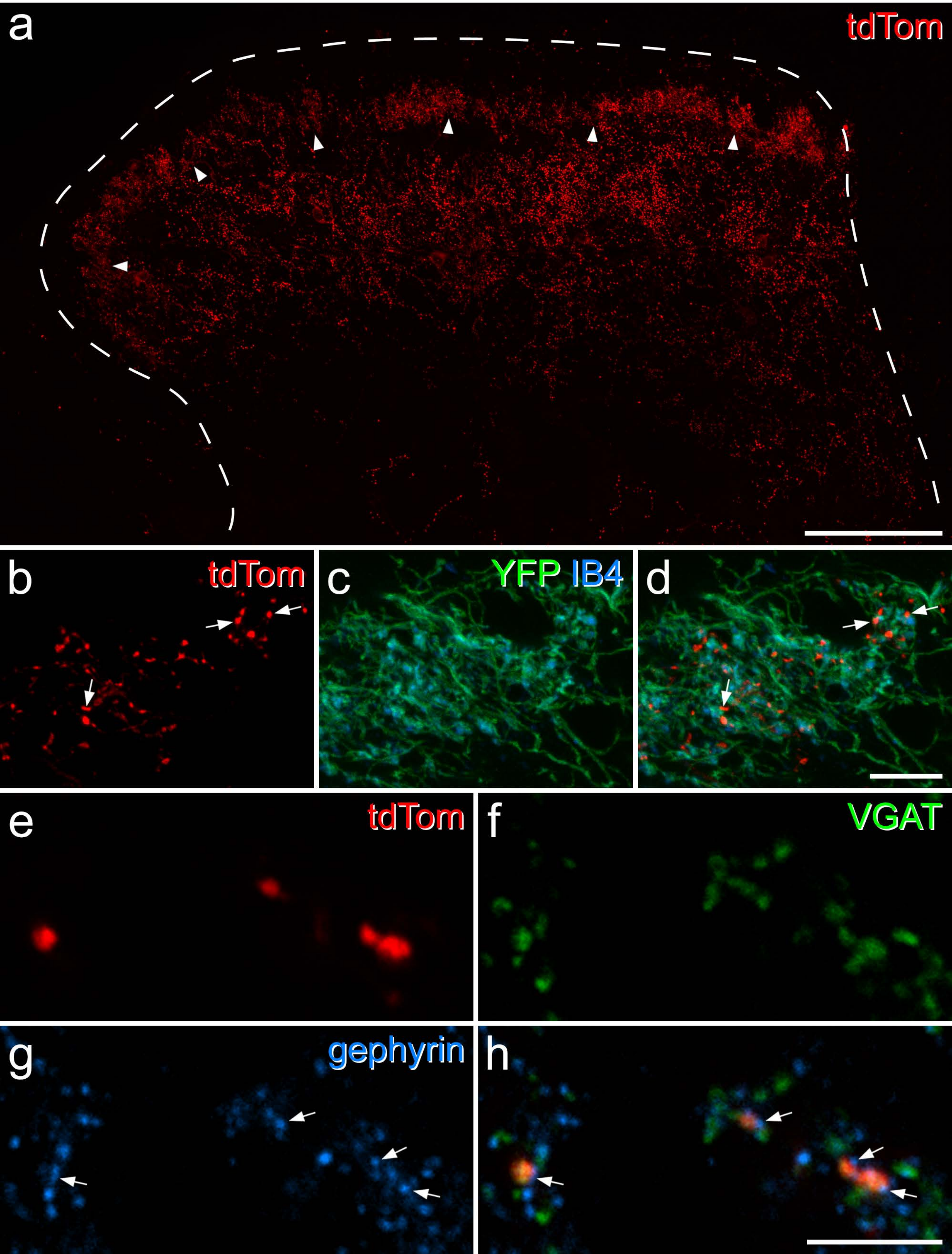

Fig S4

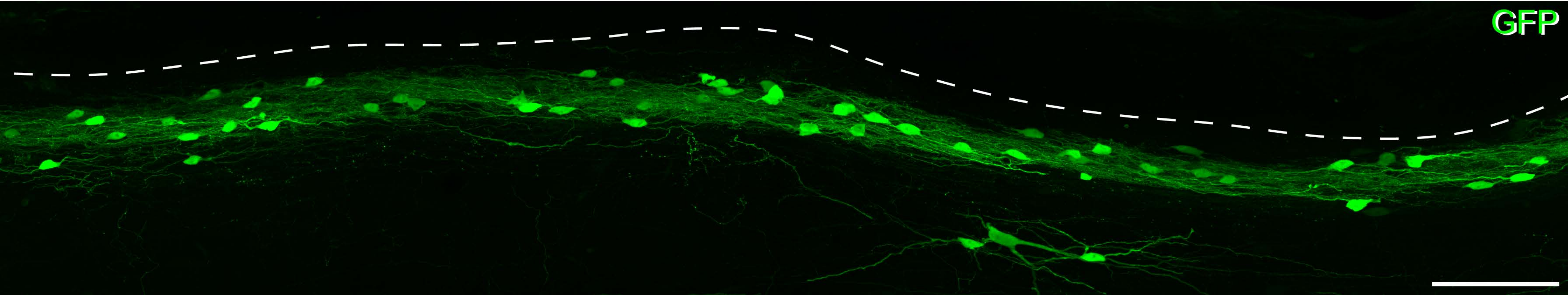

**Table S1** Mouse lines used in the study

| | Source | Reference | Catalogue number, RRID |
|---|---|---|---|
| CR::GFP | H Monyer | Caputi et al., 2009 | |
| Pnoc::GFP | HU Zeilhofer | Smith et al., 2020 | |
| $Rorb^{CreERT2}$ | DD Ginty | Abraira et al., 2017 | |
| $CR^{Cre}$ | Jackson Laboratory | | JAX:010774<br>RRID:IMSR_JAX:010774 |
| $MrgD^{ChR2\text{-}YFP}$ | Mutant Mouse Resource & Research Centers | | 036112-UNC<br>RRID:MMRRC_036112-UNC |
| $VGAT^{Flp}$ | Jackson Laboratory | | JAX:029591<br>RRID:IMSR_JAX:029591 |
| Ai9 | Jackson Laboratory | | JAX: 007909<br>RRID:IMSR_JAX:007909 |
| Ai32 | Jackson Laboratory | | JAX:012569<br>RRID:IMSR_JAX:012569 |
| Ai34 | Jackson Laboratory | | JAX:012570<br>RRID:IMSR_JAX:012570 |

**Table S2** Antibodies used in the study

| Antigen | Host species | Source | Catalogue no | Dilution | RRID |
|---|---|---|---|---|---|
| mCherry* | Rat | Invitrogen | M11217 | 1:1000 | RRID:AB_2536611 |
| mCherry* | Rabbit | Abcam | Ab167453 | 1:1000 | RRID:AB_2571870 |
| GFP** | Chicken | Abcam | ab13970 | 1:1000 | RRID:AB_300798 |
| GFP† | Rabbit | M Watanabe | | 1:10,000 | RRID:AB_2571573 |
| Pax2 | Rabbit | Life Technologies | 716000 | 1:1000 | RRID:AB_2533990 |
| Calretinin | Goat | Swant | CG1 | 1:1000 | RRID:AB_10000342 |
| Homer | Goat | M Watanabe | | 1:1000 | RRID:AB_2631104 |
| PAP | Chicken | Aves | PAP | 1:1000 | RRID:AB_2313557 |
| Somatostatin | Guinea pig | P Ciofi | | 1:1000 | |
| Gephyrin | Mouse | Synaptic Systems | 147 021 | 1:1000 | RRID:AB_2232546 |
| VGAT | Goat | M Watanabe | | 1:1000 | RRID:AB_2571623 |

*The mCherry antibody also recognises tdTomato

**The GFP antibody also recognises YFP. The chicken anti-GFP antibody was used in all cases in which YFP was to be revealed with immunofluorescence.

†The anti-rabbit GFP antibody was used for immunoperoxidase labelling in experiments involving electron microscopy.

**Table S3** RNAscope probes used in the study

| **Probe** | **Protein/peptide** | **Channel numbers** | **Catalogue numbers** |
|---|---|---|---|
| *tdTomato* | tdTomato | 2 | 317041 |
| *Tac1* | Preprotachykinin 1 | 3 | 410351 |
| *Gad1* | GAD67 | 1 | 400951 |
| *Pam* | Peptidyl-glycine alpha-amidating monooxygenase | 2 | 457271 |
| *Slc32a1* | Vesicular GABA transporter | 3 | 319191 |
| RNAscope multiplex positive control (*Polr2a*, *Ppib*, *Ubc*) | Polr2a: DNA-directed RNA polymerase II subunit RPB1; Ppib: Peptidyl-prolyl cis-trans isomerase B; Ubc: Polyubiquitin-C | 1,2,3 | 320881 |
| RNAscope multiplex negative control (*dapB*) | dapB: 4-hydroxy-tetrahydrodipicolinate reductase (derived from B Subtilis) | 1,2,3 | 320871 |

### Supplementary figure legends

Fig S1 Labelling of iCRs in the Pnoc::GFP mouse. **a**-**c**: a transverse section from one of these mice has been reacted to reveal GFP (green), Pax2 (blue) and calretinin (CR, red). **d**: a merged image. The inset images show 3 GFP-labelled cells (arrows), which are immunoreactive for both Pax2 and calretinin. The box in **d** shows the area from which the inset was taken. Main images are maximum intensity projections of 15 confocal optical sections at 2 μm z-spacing. The inset images are from a single optical section. Scale bars = 200 μm (main images) and 25 μm (insets).

Fig S2 Fluorescent *in situ* hybridisation for *Tac1*, *Slc32a1*, and *Pam* mRNAs in lamina II of a wild-type mouse. *Slc32a1* encodes the vesicular GABA transporter, and is present in inhibitory interneurons, while *Pam* codes for peptidyl-glycine alpha-amidating monooxygenase, the enzyme that amidates substance P. **a**-**c** show individual channels, while **d** is a merged image that also includes nuclear staining with NucBlue (grey). This field contains several *Tac1*-containing cells, four of which also contain *Slc32a1* and are therefore inhibitory (arrowheads). Two excitatory *Tac1*-containing cells (identified by lack of *Slc32a1*) are marked with arrows. The density of transcripts for *Pam* is much lower in the four inhibitory Tac1 cells than in the two excitatory cells. Images are maximum intensity projections of confocal optical sections (1 μm z-separation) through the full thickness of the section. Scale bar = 10 μm.

Fig S3 Axons of Rorb cells revealed by staining for tdTomato (tdTom) in the $Rorb^{CreERT2}$;Ai34 cross. **a**: a scan through the dorsal horn of a $Rorb^{CreERT2}$;Ai34;$MrgD^{ChR2\text{-}YFP}$ mouse reveals a dense band of tdTomato labelling in lamina II (marked by arrowheads), as well as staining in deeper laminae (III-IV). There is a partial separation between these two layers. **b-d**: a section from the same cross scanned through lamina II to reveal tdTom (red), YFP (green) and IB4 binding (blue). The tdTomato-labelled boutons in lamina II are closely associated with MrgD axonal boutons, revealed by IB4 binding in YFP-labelled varicosities. Arrows point to appositions between individual tdTomato-labelled and MrgD+/IB4+ boutons. **e-h**: a scan from lamina II of a $Rorb^{CreERT2}$;Ai34 mouse, immunostained to reveal tdTom (red), VGAT (green) and gephyrin (blue). Four tdTomato-labelled boutons are present in this field, and all are VGAT-immunoreactive. Each of these is in contact with a gephyrin punctum (arrows in **g** and **h**). The image in **a** is a projection of 27 optical sections at 1 μm z-separation, those in **b-d** are projections of 11 optical sections at 0.3 μm z-separation, while those in **e-h** are from a single optical section. Scale bars: 100 μm (**a**), 10 μm (**b-d**) and 5 μm (**e-h**).

Fig S4 Intersectional targeting of inhibitory Tac1 cells. A sagittal section through the region of the injection site in a $Tac1^{Cre}$;$VGAT^{Flp}$ mouse that had received an injection of AAV.$C^{on}$/$F^{on}$.GFP, immunostained to reveal GFP. The dashed line represents the dorsal surface of the grey matter. Note that the great majority of GFP labelled cells are located in lamina II, and that their processes remain in this lamina. The image is a projection of 27 confocal optical sections at 1 μm z-spacing. Scale bar = 100 μm.
